# Macrophages utilize a bet-hedging strategy for antimicrobial activity in phagolysosomal acidification

**DOI:** 10.1101/470617

**Authors:** Quigly Dragotakes, Kaitlin Stouffer, Man Shun Fu, Yehonatan Sella, Christine Youn, Insun Yoon, Carlos M. De Leon-Rodriguez, Joudeh B. Freij, Aviv Bergman, Arturo Casadevall

## Abstract

Microbial ingestion by a macrophage results in the formation of an acidic phagolysosome but the host cell has no information on the pH susceptibility of the ingested organism. This poses a problem for the macrophage and raises the fundamental question of how the phagocytic cell optimizes the acidification process to prevail. We analyzed the dynamical distribution of phagolysosomal pH in murine and human macrophages that had ingested live or dead *Cryptococcus neoformans* cells, or inert beads. Phagolysosomal acidification produced a range of pH values that approximated normal distributions, but these differed from normality depending on ingested particle type. Analysis of the increments of pH reduction revealed no forbidden ordinal patterns, implying that phagosomal acidification process was a stochastic dynamical system. Using simulation modeling, we determined that by stochastically acidifying a phagolysosome to a pH within the observed distribution, macrophages sacrificed a small amount of overall fitness to gain the benefit of reduced variation in fitness. Hence, chance in the final phagosomal pH introduces unpredictability to the outcome of the macrophage-microbe, which implies a bet-hedging strategy that benefits the macrophage. While bet hedging is common in biological systems at the organism level, our results show its use at the organelle and cellular level.

## Introduction

### *Audaces fortuna iuvat* (Fortune favors the bold) - Virgil

Phagocytosis is a fundamental cellular process used by unicellular organisms for nutrient acquisition as well as by host immune cells for microbial defense. The parallels between food acquisition and immunity have led to the suggestion that these two processes had a common evolutionary origin^1^. The process of phagocytosis results in the formation of a phagolysosome, a dynamic membrane bounded organelle, which represents a critical location in the struggle between the host and ingested microbial cells^2^. Microbial ingestion into phagosomes results in exposure to host cell microbicidal mechanisms, which leads to death for some microbes while others survive by subverting critical aspects of phagosome maturation and by damaging phagolysosome structural integrity.

The process of phagosomal maturation, encompassed by the fusion of the phagosome with lysosomes and lumen acidification, is a complex choreography that includes the recruitment of V-ATPase from lysosomes to the phagolysosome^2,3^ and many other protein components^4^. Proton pumping into phagolysosomal lumen results in acidification that inhibits microbes and activates antimicrobial processes that are microbicidal. Consequently, some types of microbes, such as *Mycobacterium tuberculosis* and *Histoplasma capsulatum*, interfere with phagosomal maturation and acidification to promote their intracellular survival. The extent of phagosomal acidification is determined by numerous mechanisms that include proton flux through the pump, proton consumption in the phagosomal lumen, and backflow into the cytoplasm^5^. Phagosome acidification in macrophages is rapid with a pH of 6 being reached within 10 min after ingestion^6^ and 5.4 by 15-20 min^7^.

*Cryptococcus neoformans* is a facultative intracellular pathogen^8^. Upon ingestion by macrophages, *C. neoformans* resides in a mature acidic phagolysosome^9^. The outcome of macrophage-*C. neoformans* interaction is highly variable depending on whether the fungal cell is killed, inhibited, or unaltered. If not killed, *C. neoformans* can replicate intracellularly, resulting in variable outcomes which include death and lysis of the host cell, non-lytic exocytosis^10,11^, transfer to another macrophage^12,13^, or phagosomal persistence. A critical variable in determining the outcome of the *C. neoformans*-macrophage interaction is phagosomal membrane integrity, with maintenance of this barrier conducive to control of intracellular infection while loss of integrity leads to host cell death^14^.

Prior studies of *C. neoformans* phagosomal acidification measured great variation in the pH of individual phagolysosomes^14–16^. The pH of cryptococcal phagolysosomes is affected by several microbial variables that include urease expression^15^, phagosomal membrane integrity^14^, and the presence of the cryptococcal capsule with glucuronic acid residues that can influence the final pH via acid base properties^17^. In addition, the capsule of *C. neoformans* increases in diameter as part of a stress response which can potentially affect the phagolysosomal pH through increasing the phagolysosome volume, thus diluting its contents and promoting membrane damage through physical stress^14^.

In this study, we analyzed the time dependence of the phagolysosomal pH distribution in murine and human macrophages and conclude that acidification is a stochastic process in which macrophages default to a range of pHs with random variation. We demonstrate that, in doing so, macrophages employ a *bet-hedging* strategy by sacrificing a small amount of overall fitness to ensure broad survivability. Thus, we suggest that chance, in the form of stochastic dynamics, could be an important strategy in phagosomal acidification, which benefits the macrophage by bet hedging against a range of pathogens, and could echo through the immune process to introduce a fundamental uncertainty in the outcome of microbe-macrophage interactions.

## Results

We previously reported a wide distribution of phagolysosomal pH after the ingestion of *C. neoformans* by murine bone marrow derived macrophages (BMDMs)^14,15^. Given that the growth rate of *C. neoformans* is highly affected by pH^15,18^ and that the outcome of the *C. neoformans*-macrophage interaction is determined in the phagolysosome^14,19,20^, we decided to analyze the distribution of phagolysosomal pH mathematically to gain insight into the dynamics of the acidification process. Understanding the fundamental process of phagolysosomal acidification, and if there are overarching rules or determinism involved, could provide key insights to disease pathology of any pathogen, which would interact with the phagolysosome. A scheme of the method used to determine phagosomal acidification with representative data from polystyrene bead phagocytosis experiments are shown in Figure 1. To determine the pH of a phagolysosome, particles are opsonized with an Oregon Green conjugated Ab. Oregon Green is pH insensitive at Ex440/Em520 and pH sensitive at Ex488/Em520, allowing pH to be measured by quantifying the ratio of fluorescent intensity between each wavelength. We were able to establish robust standard curves from known pH controls and calculate unknown phagolysosome measurements (Figure 1B).

**Figure 1.**
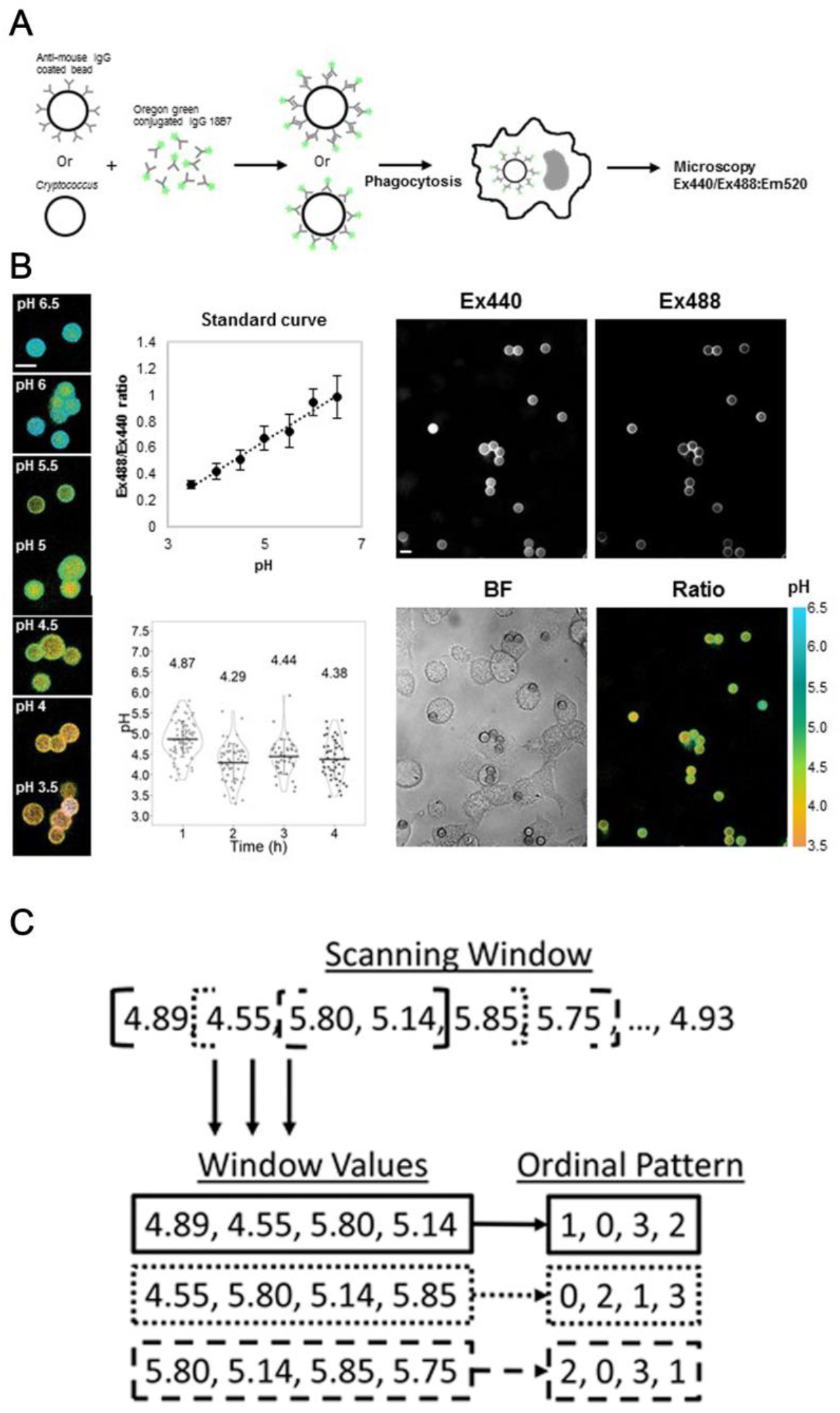
Experimental designs for described methods. **A.** Experimental design for phagolysosomal pH measurement. Particles are opsonized with mAb 18B7 conjugated to fluorophore OregonGreen. **B.** Measurement of phagolysosomal pH. Measurements are based on ratiometric measuring of Ex440/Em520 and Ex488/Em520. **C.** Experimental design for the detection of Chaos within a system. A scanning window generates sets of windows values. Ordinal patterns are then generated by ordering the terms in each scanning window from least to greatest.

### Murine macrophage phagolysosomes acidify stochastically

To determine whether phagolysosomal acidification is a deterministic or stochastic dynamical process, we employed a permutation spectrum test^21^ in which the distribution of ordinal patterns occurring in subsets of our full dataset was analyzed (Figure 1C). Ordinal patterns simply refer to the order of each measurement in terms of value, in our case the phagolysosomal pH, within a scanning window, which parses all measurements within each condition, exemplified in Figure 1^22–25^. Here we found a 4-unit window size to be the most appropriate. We found no forbidden patterns at any time evaluated for any of the pH distributions resulting from the synchronized ingestion of beads, alive *C. neoformans,* or heat-killed *C. neoformans*, which implies that the acidification is a stochastic process (Figure 2). A forbidden ordinal is an ordinal pattern that does not appear during the time frame of our experiment. Despite the overall population of macrophages showing chaotic dynamics, we considered whether the Chaotic signatures occurred at the individual cellular level. That is, whether slight differences at the initiation of phagocytosis propagated through maturation and acidification resulting in two phagolysosomes within the same cell with different acidifications. To determine if this was the case, we analyzed the phagosomal pH in pairs of bead containing phagolysosomes for which each pair was within a single macrophage. We chose macrophages in which the ingested beads were visibly separated. We found that the differences between phagolysosomal pH measurements of this cohort of phagolysosomes yields a normal distribution centered at 0 with non-zero values, suggesting each individual phagosome has an independent target phagolysosomal pH (SFig 1). The ordinal pattern analysis was repeated on several other strains and conditions throughout the project and at no point did we observe forbidden ordinal patterns (SFig 2A), again suggesting the process exhibits no signature of deterministic chaos^22–25^. Given that beads are inert and cannot modify pH, we reasoned that their corresponding phagolysosomes would be the closest approximation of a default acidification state and would therefore represent a “baseline” to which we could compare phagolysosome acidification dynamics in other conditions.

**Figure 2.**
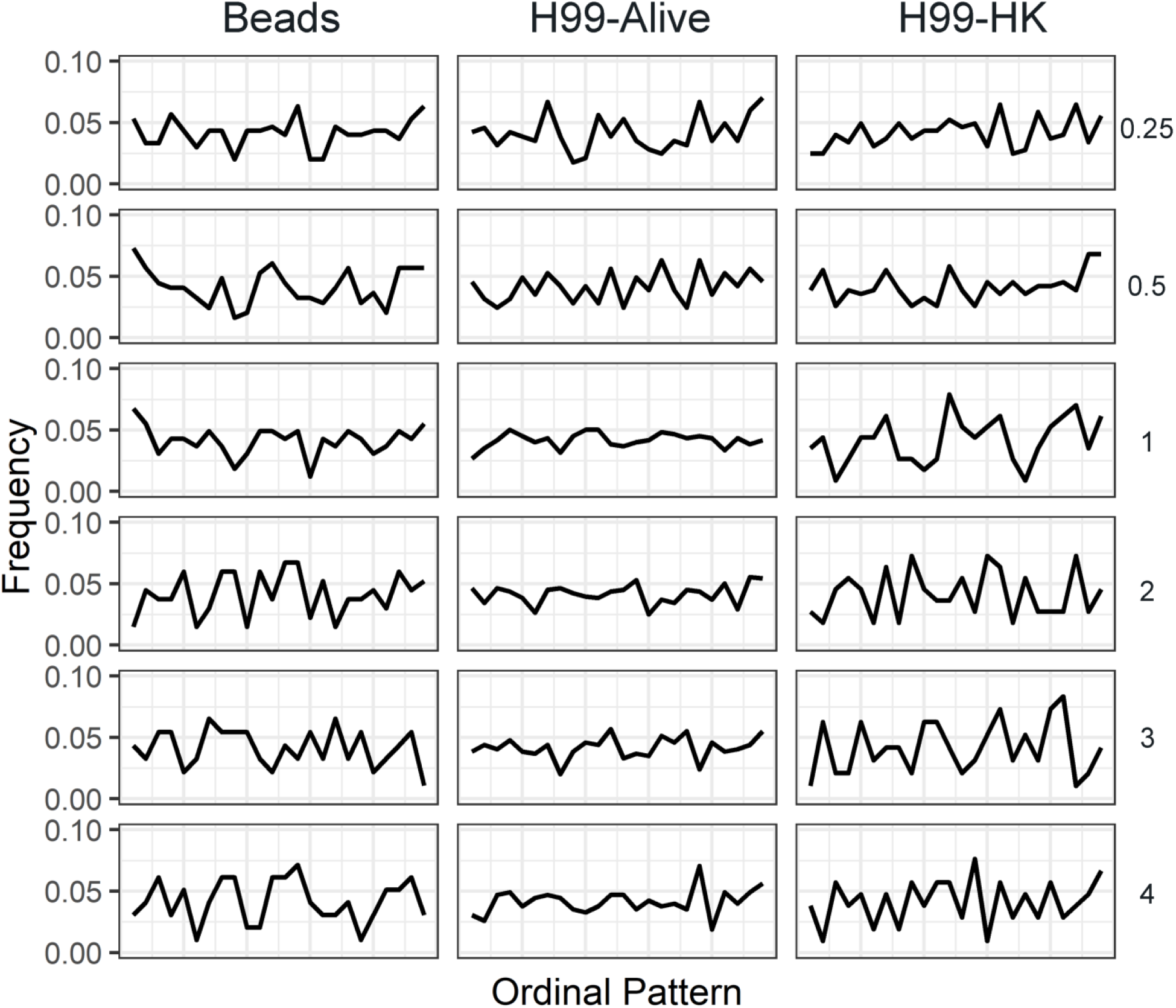
Murine macrophage phagolysosomes acidify stochastically. The frequency of each 4-unit ordinal pattern for each ingested particle: beads, live *C. neoformans*, and dead *C. neoformans* at various HPI. Note that all ordinal patterns have a non-zero frequency.

### Bead ingested murine macrophage phagolysosomes stabilize to a normally distributed pH

To probe the dynamics of the phagolysosomal acidification system at this baseline, we analyzed several hundred individual phagolysosomal pH measurements at various time intervals after BMDMs had ingested inert beads (Figure 3A). To determine whether phagolysosome pH measurements followed a normal distribution, the measured relative pH values were fit to a predicted normal distribution using the “fitdistrplus” R statistical package then analyzed via Q-Q plots and the Shapiro-Wilk normality test. We found that phagolysosome pH value distributions did not approximate normality at early times (15 and 30 min), but mostly did at intervals of and past 1 h post infection. We were able to reject the Shapiro-Wilk null hypothesis only at 15 min, 30 min, and 3 h (Figure 3B). We hypothesized that not all phagolysosomes were fully mature before 1 h and reasoned that the bimodal appearance of pH at early time intervals is likely due to the population of phagosomes being at different stages of maturity. Interestingly, the pH distribution for the phagolysosomes containing live or dead *C. neoformans* did not approximate normality even at later times. We rejected Shapiro-Wilk normality for each of these samples except for dead *C. neoformans* at 3 h. Q-Q plots supported the notion that each sample skews further from normality compared to bead ingested macrophage phagolysosome pH distributions (SFig 3). Considering that macrophages default to a random pH from a particular normal distribution and that *C. neoformans* invest in disrupting this process, we hypothesized that this system must confer some host benefit. Thus, we decided to compare the pH distribution of mature bead ingested phagolysosomes to a range of pHs tolerated by relevant potential pathogens.

**Figure 3.**
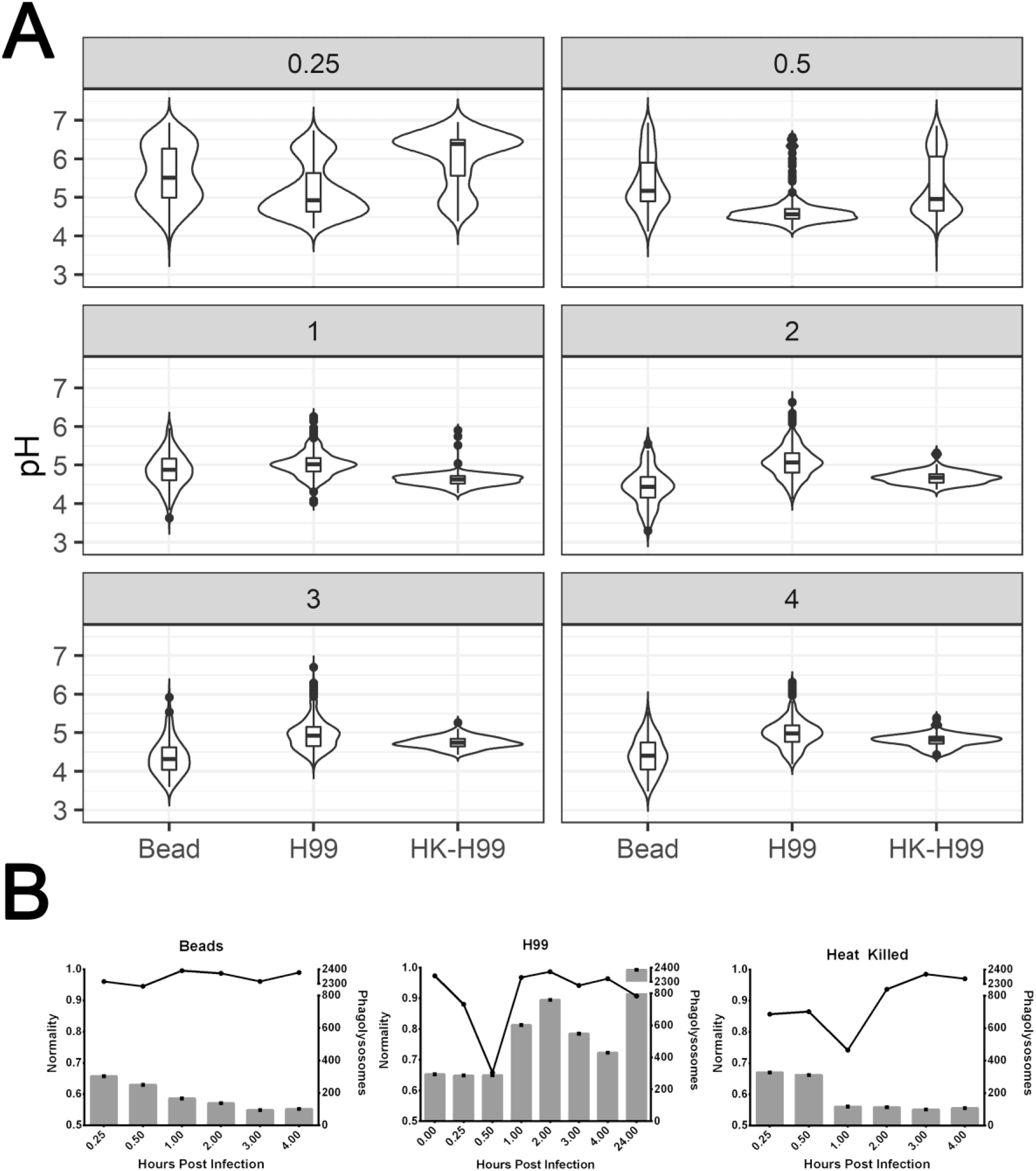
Bead ingested murine macrophage phagolysosomes stabilize to a normally distributed pH. **A.** Distributions of measured phagolysosomal pH after ingestion of various particles by BMDMs at various hours post infection (gray box value) for bead, live *C. neoformans* (H99), and dead *C. neoformans* (HK-H99). **B.** Visualization of estimated normality via Shapiro-Wilk (line) and total phagolysosome count (bar) for each sample.

### Acidification Dynamics are Closely Related to Maturation

To probe whether the observed normality and stochasticity is a result of dynamics in acidification or in the maturation process, we analyzed the intensity of two phagolysosomal maturation markers, EEA1 and V-ATPase, over the same time course. Using relative intensity of each immunofluorescent staining as a surrogate measurement of maturation, we found that, again, no samples had forbidden ordinal patterns, suggesting phagolysosomal maturation, when measured by the accumulation of these markers, is also stochastic in nature (SFig 4). However, neither of the fluorescence intensity measurements from these markers approximated normality at any time, with sufficient skewing to reject the Shapiro Wilk null hypothesis (SFig 5). Skewing of these measurements away from normality could reflect cytoplasmic speckling, limitations of fluorescent microscopy resolution, non-linearity of fluorescence signals and the inherent complexity of such a system, such that we are not confident to reject the notion that these processes indeed demonstrate a normal distribution. We hypothesized that macrophages may experience limited resources in terms of the number of available V-ATPase pumps at any given time and that we might see a correlation between intensity of VATP staining around individual phagosomes in cells that had ingested one versus multiple beads. We found no evidence of a correlation between V-ATPase staining, or EEA1 for that matter, and total number of ingested particles (SFig 6).

### BMDM phagolysosomes acidify to a pH range suboptimal for growth of soil and pathogenic microbes

The pH of soils varies greatly from acidic to alkaline based on a variety of conditions that, in turn, determine the associated microbiome^26^. Soils contain many pathogenic microbes including *C. neoformans*. Since the phagolysosome is an acidic environment, we reasoned that microbes that thrive in acidic soils could proxy for the types of microbes that hosts, and thus macrophages, could encounter, and pose a threat to the cell/host due to their acidophilic nature (ex. *Cryptococcus neoformans*). Hence, we compared^27,28^ the distribution of pH values from mature phagolysosomes (HPI ≥ 1 h) which had ingested latex beads, as a measure of the range of acidities generated in the absence of microbial modulation, relative to published soil microbe growth data as a function of pH (Figure 4). The latex bead pH distribution is narrow and centered at a pH of about 4.5, which corresponds to a pH that significantly reduces the optimal growth even for microbes in acidic soils. To generate a more relevant comparison to human disease models, we also searched out the pH tolerance of 27 significant human pathogens which macrophages are likely to encounter during human infectious diseases (Supplemental Table I). We then compared these distributions to the pH distribution in bead ingested macrophage phagolysosomes (Figure 4). The distribution of this default macrophage pH heavily overlaps with the more inhibitory pH regions when compared to both soil and known pathogen pH tolerances. Taken together these data suggest that the low phagosomal pH is itself a defense mechanism, with a distribution that manifests bet hedging by defaulting to a range, rather than single value, inhibitory to most of the spectrum of pathogens the macrophage is likely to encounter.

**Figure 4.**
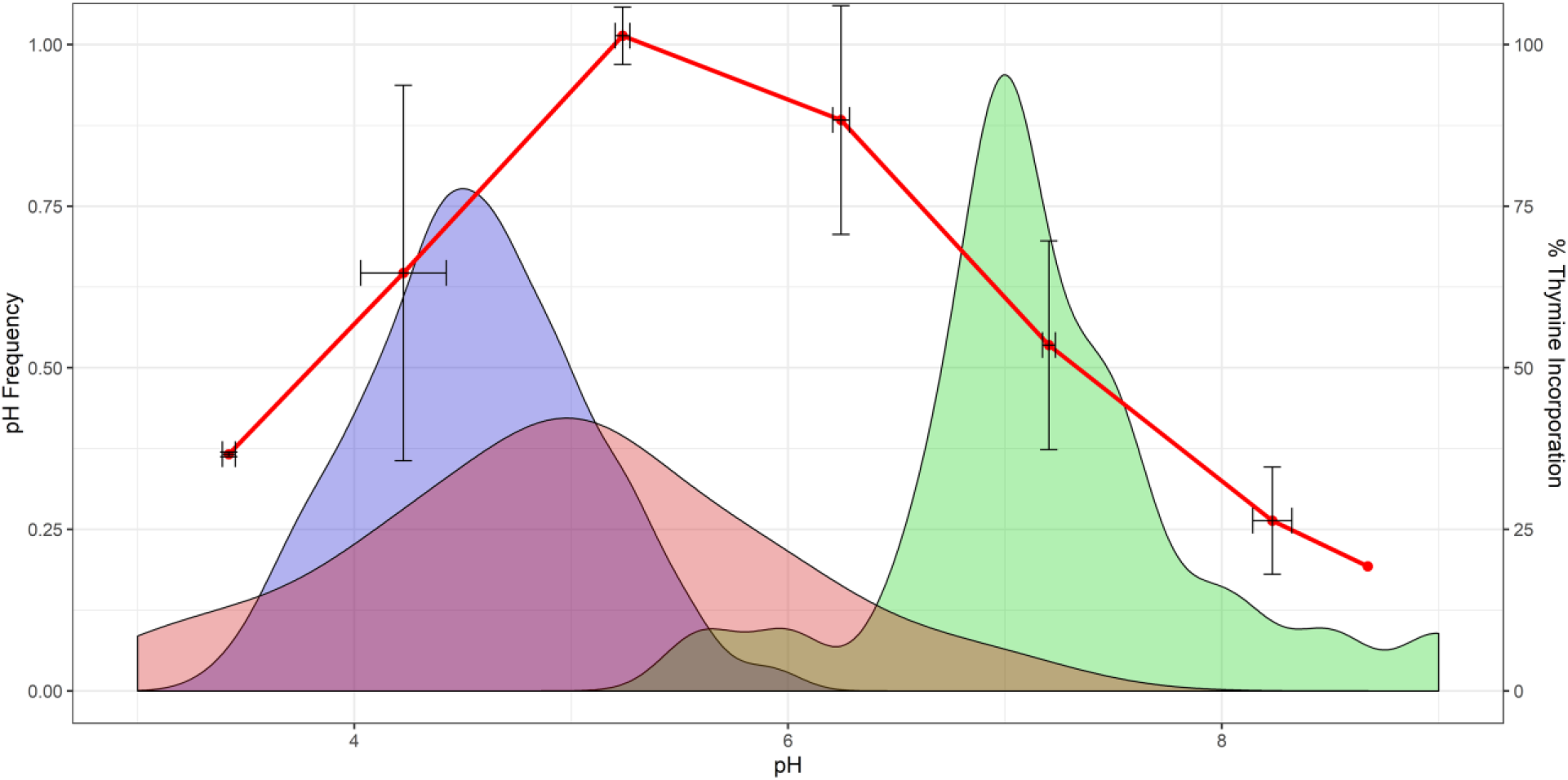
BMDM phagolysosomes acidify to a default pH range suboptimal for soil microbe and pathogen growth. Growth rates of soil bacteria^27,28^ as percent thymine incorporation (red line) and distribution of minimal culturable pH for 27 human pathogens (red fill) compared to the distribution of observed phagolysosomal pH after bead ingestion and phagolysosome maturation, 1-4 HPI (blue fill). The bead pH distribution overlaps unfavorably with minimal growth conditions of microbes, and only minimally overlaps with the optimal growth pH range of the same 27 pathogens (green fill).

### Simulations of macrophage populations show stochastic pH as a bet-hedging strategy

Bet hedging is generally understood as a strategy whereby an organism decreases variation in fitness at the expense of a small decrease in mean fitness. To test whether variation in macrophage phagolysosome pH constitutes a diversified bet-hedging strategy, we first modeled host survival rate as a function of final phagolysosomal pH in the context of a pathogen randomly selected from Supplemental Table I. This analysis modeled phagolysosomal pH as a normal distribution with mean μ and standard deviation σ varied over a range of possible values in order to test how bet hedging depends on these parameters. To each phagolysosomal pH, we associated a fitness value which models survival likelihood against pathogens, using the list of pathogens (Table 1), with their viable pH ranges collected from the literature, and we computed the distribution of fitness (p) as phagolysosomal pH varies (Figure 5A,B). As expected in bet hedging, we observed decreases in standard deviation of fitness with increasing σ of phagolysosomal pH (Figure 5C). Mean fitness was mostly unchanged with changing standard deviation, such that even in the most extremes deviations in phagolysosomal pH there were only slight changes in fitness in either direction.

**Figure 5.**
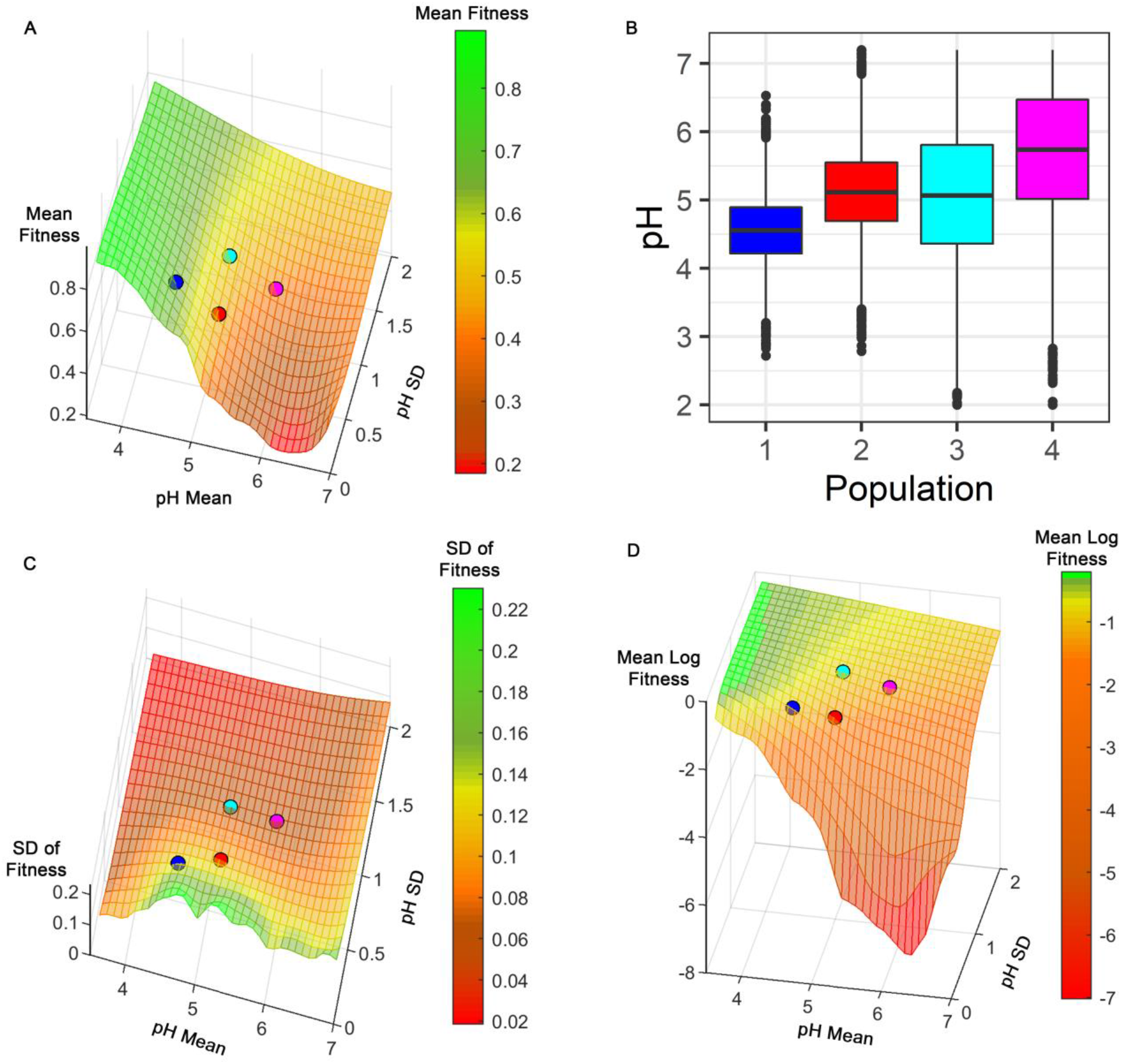
Simulated macrophage populations show stochastic pH as bet-hedging strategy. Success of host-microbe interactions are visualized as macrophage populations faced with randomly selected pathogens and acidified to random pHs from normal distributions of mean μ and standard deviation σ. Meshes represent host macrophage fitness, deviation in fitness, and log mean fitness. Plotted points represent measured data of H99 containing murine (red) and human (magenta) or bead containing murine (blue) and human (cyan) phagolysosomes. **A.** Mean macrophage survival (Z axis, colorbar) increases significantly as pH lowers, and mostly unchanged with standard deviation. **B.** Examples of simulated populations based on colored points in panel A. Each combination of mean and SD from the axes of panel A represent a unique population of macrophages with a fitness represented by the Z axis and colored mesh. **C.** Deviation in host fitness (Z axis, colorbar) dramatically decreases by increasing standard deviation of pH, and mostly unaffected by shifts in mean pH. **D.** Logarithmic measurement of host fitness (Z axis, colorbar) to observe long term trends applicable to a bet-hedging strategy.

Another way to formalize the emergence of bet hedging as an evolutionary strategy is by considering the average long-term rate of growth. Thus, we considered a multiplicative model, in which the growth rate (*r*_*T*_) over a span of time (T) is defined as the geometric mean of *ρ*_1_, …, *ρ*_*T*_, where for each t ranging from 1 to T, *ρ*_*t*_ is the fitness at time t. Taking logs, log(*r*_*T*_) is the average of log(*ρ*_1_),…,log(*ρ*_*T*_). By the law of large numbers, if we assume each *ρ*_*t*_ is independent and identically distributed, this average will approach E(log(ρ)). Thus E(log(ρ)) is the operative quantity to be maximized, rather than E(ρ). Note that applying the log transformation has the effect of placing heavier penalties on fitness values that are close to 0, so that maximizing mean log fitness will tend to encourage lower standard deviation in fitness^29^. Thus, increased mean log fitness is another indication of bet-hedging which directly relates to long-term growth. We indeed observe increased mean log fitness with increasing σ of phagolysosomal pH (Figure 5D).

To probe whether our simulations would reflect biological responses, we analyzed macrophage phagolysosomal pH in which cells treated with Chloroquine, a weak base that localizes to the phagolysosome. According to our model the increased shift in mean pH from chloroquine would result in a lower overall mean log fitness as a result of shifting the mean pH closer to 6, a more tolerable region for most of the candidate pathogens. Thus, we compared previously reported data^9^ of *C. neoformans* containing phagolysosomes treated with chloroquine to our data from the respective time interval and found that chloroquine treatment results in a drastically reduced overall mean log fitness (−5.323) compared to the mean log fitness of our non-chloroquine treated data of the respective condition (−2.486).

### Time intervals from ingestion of *C. neoformans* to initial budding are stochastic

*C. neoformans* replication rate is highly dependent on pH^15^. Consequently, we hypothesized that if phagolysosomal acidification followed stochastic dynamics, this would be reflected on the time interval from ingestion to initial replication. Analysis of time intervals to initial fungal cell budding events revealed stochastic dynamics with no evidence of forbidden ordinal patterns (SFig 2B). Similar results were observed for initial budding of wild type and urease negative strains of *C. neoformans*, which reside in phagolysosomes that differ in final pH as a result of ammonia generation from urea hydrolysis. Acidification intervals for both strains were stochastic, despite the fact that phagolysosomes of urease deficient strains are approximately 0.5 pH units lower than those of wild type strains^15^.

### Trained murine macrophages have inverse acidification dynamics

Trained immunity has recently been shown to influence repeated infection in monocyte populations not exposed to the adaptive immune system^30^. To determine whether initial exposure to a pathogen has an effect in the dynamics of this system we exposed BMDMs to *C. neoformans,* resolved the infection with antifungals, and measured phagolysosome acidification dynamics upon reinfection. We found that pH distribution of phagolysosomes from macrophages previously trained as described also exhibited stochastic behavior (Figure 6A), veering away from a normal distribution of pH (Figure 6B and SFig 8). However, trained BMDMs on average exhibited a significantly lower initial pH, which became significantly higher compared to untrained BMDMs over time (Figure 6C). Fully understanding this system will require significant further study outside the scope of this manuscript. In this regard we note that amphotericin B is a powerful activator of macrophages^31^ and that *C. neoformans* residence inside macrophages is associated with host cell damage^32,33^. Hence, the effects we observe could be the aggregate of several influences in the system. Nevertheless, there is a clear suggestion of a historical effect on which pH distribution a macrophage will employ. This may also suggest an adaptive component to the macrophage bet-hedging strategy.

**Figure 6.**
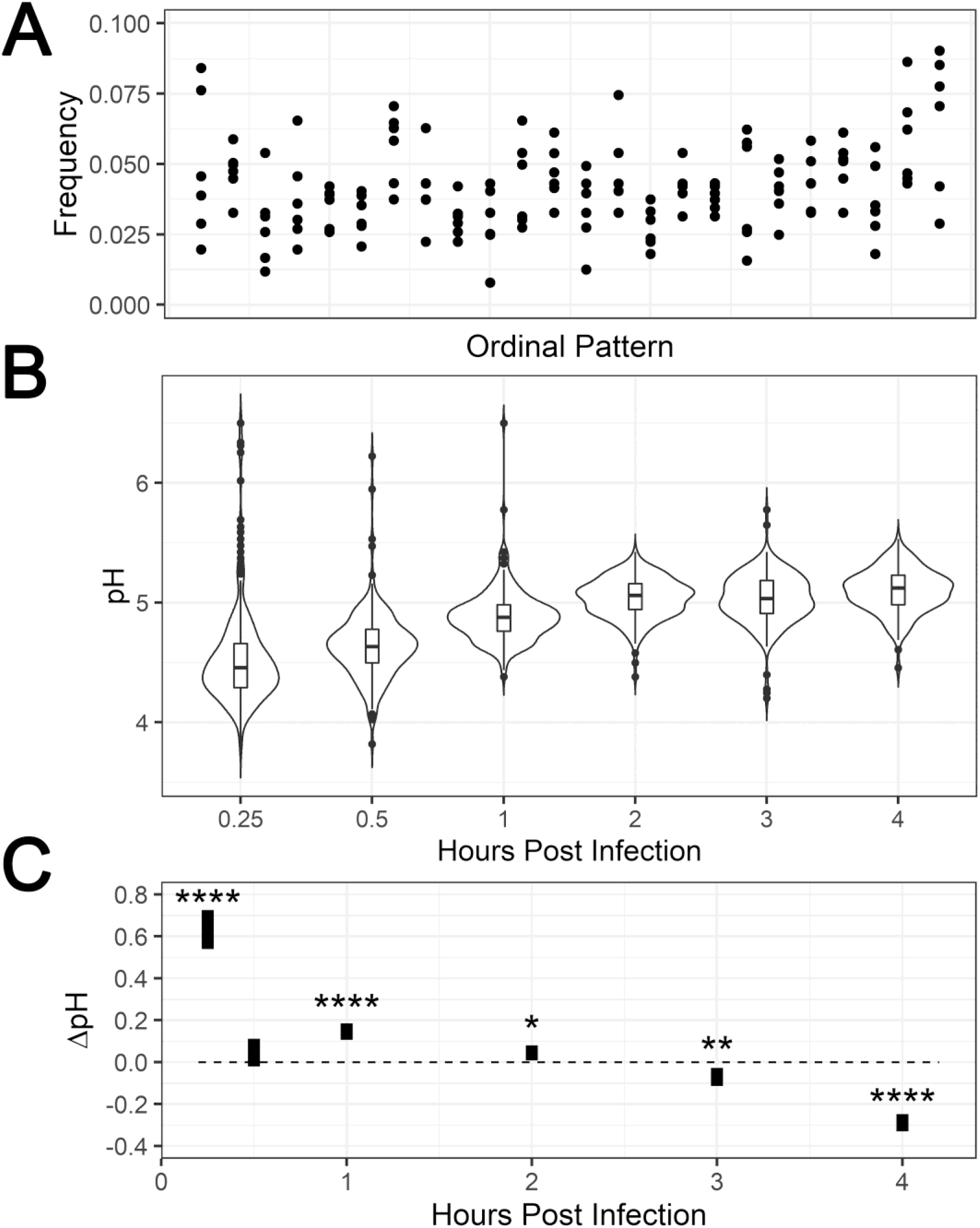
Trained murine macrophages have inverse acidification dynamics. **A.** Ordinal pattern analysis of phagolysosome pH in trained BMDMs. All ordinal patterns appear at a non-zero frequency. **B.** pH distributions of phagolysosomes at various HPI. **C.** Difference (95% CI) after subtracting mean trained phagolysosome pH from mean untrained phagolysosome pH at each timepoint. *, **, **** denote *P* < 0.05, 0.01, and 0.0001 respectively via two tailed t-test.

### Differently polarized macrophages acidify stochastically but not using this bet hedging system

To probe whether M0 and M2 polarized macrophages acidify with the same dynamics of M1 macrophages, we repeated these experiments with macrophages that were either not stimulated, or stimulated with IL-4 to skew towards M2. First, we found that regardless of the polarization skew all macrophages acidified stochastically (Figure 7A). Second, we found that the phagolysosomal pH distributions differed overall with M2 macrophages having the highest mean pH, followed by M0 and then followed by M1 skewed (Figure 7B). Most striking was the observation that M0 and M2 skewed macrophages did not manifest a normally distributed pH range as observed with M1 skewed macrophages. Instead, M0 macrophages consistently yield a bimodal distribution even after 1 h. The M2 macrophages phagosomal pH distribution started with a heavy tail of higher pH and eventually stabilize to a bimodal distribution. These data suggest that while the macrophages have the same underlying acidification dynamics, they do not share the betting strategy of M1 skewed macrophages. Thus, we estimated each population of macrophages likelihood to survive when faced with the same list of human pathogens modeled after bimodal distributions estimated from the observed data (SFig 9). After comparing these simulations, we found that M1 macrophages by far have the highest mean log fitness, followed by M0, followed by M2 (Figure 7C). Our model shows M1 macrophages as resistant to these infectious agents with M2 permissive.

**Figure 7.**
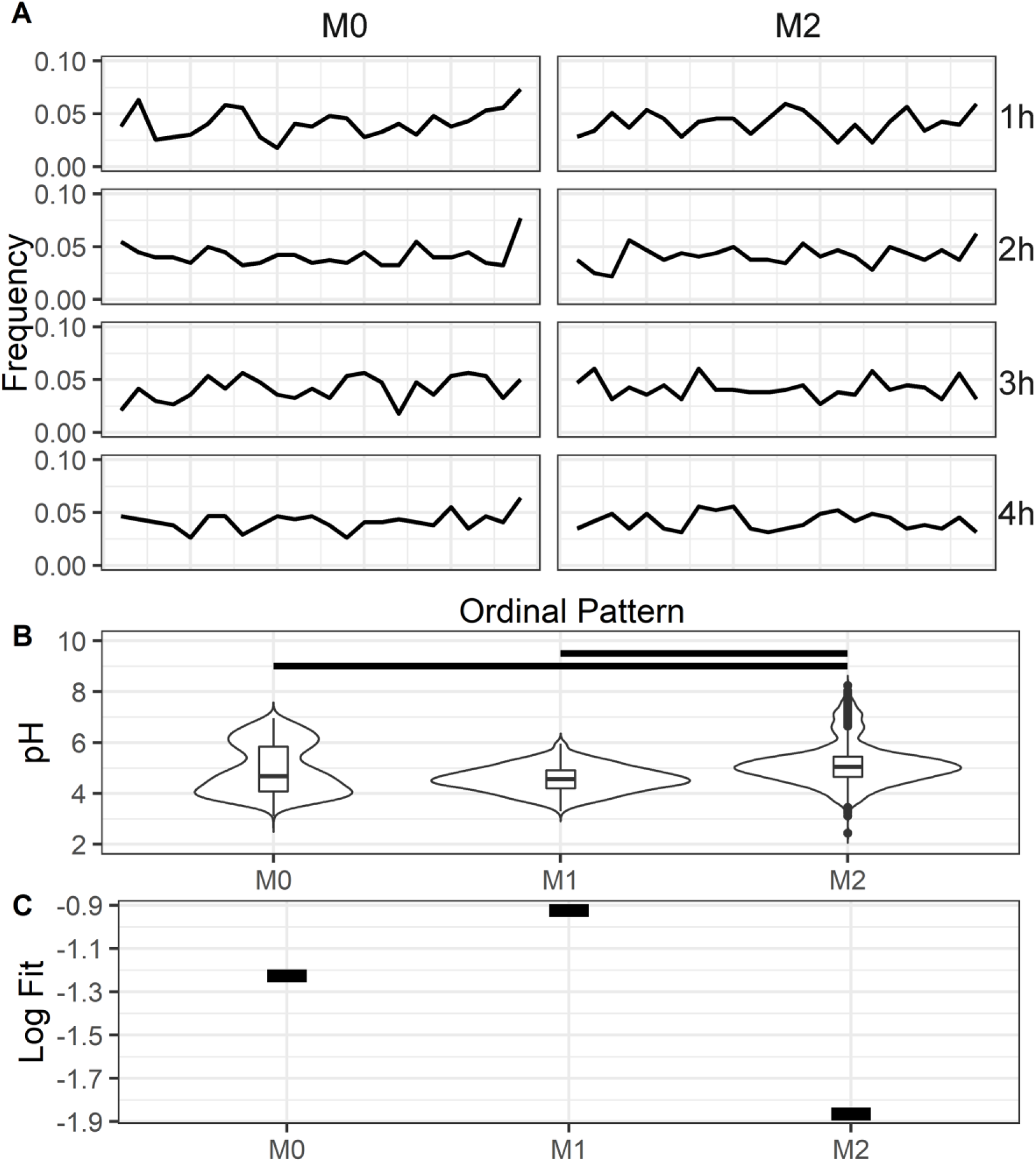
Phagolysosome dynamics of macrophages skewed toward different polarization states. **A.** Ordinal patterns of bead containing phagolysosomes at various HPI. **B.** Bead containing phagolysosome pH distributions of differently polarized macrophages. Black bars represent *P* < 0.0001 via Kruskal-Wallis test with Wilcox rank pairing test. **C.** Mean log fitness of bead containing phagolysosomes according to our bet hedging model.

### Human monocytes acidify stochastically and approximate normality

To determine how closely the murine system resembled human acidification dynamics, we isolated macrophages from human peripheral blood monocytes and repeated these experiments with beads and live *C. neoformans*. We found that acidification intervals in human cells were also stochastic in nature (Figure 8A). Additionally, human cells that ingested inert beads were normally distributed at the 15 min and 1 h time intervals. Even though the times skewed away from normality, the skew was not as severe as that observed in yeast containing phagolysosomes (Figure 8B). We hypothesize that some of this skewing could result from different dynamics due to the different background inherent to human donors, which differ from the mouse system in which cells are isolated from genetically identical individuals (SFig 10). Furthermore, within the context of our simulation, if we model the phagolysosome pH values of a population of human macrophages as a normal distribution with mean and standard deviation determined from pH values observed across all time points (5.08 and 1.08, respectively), the resulting mean log fitness is high with dramatically reduced deviation in fitness (Figure 5).

**Figure 8.**
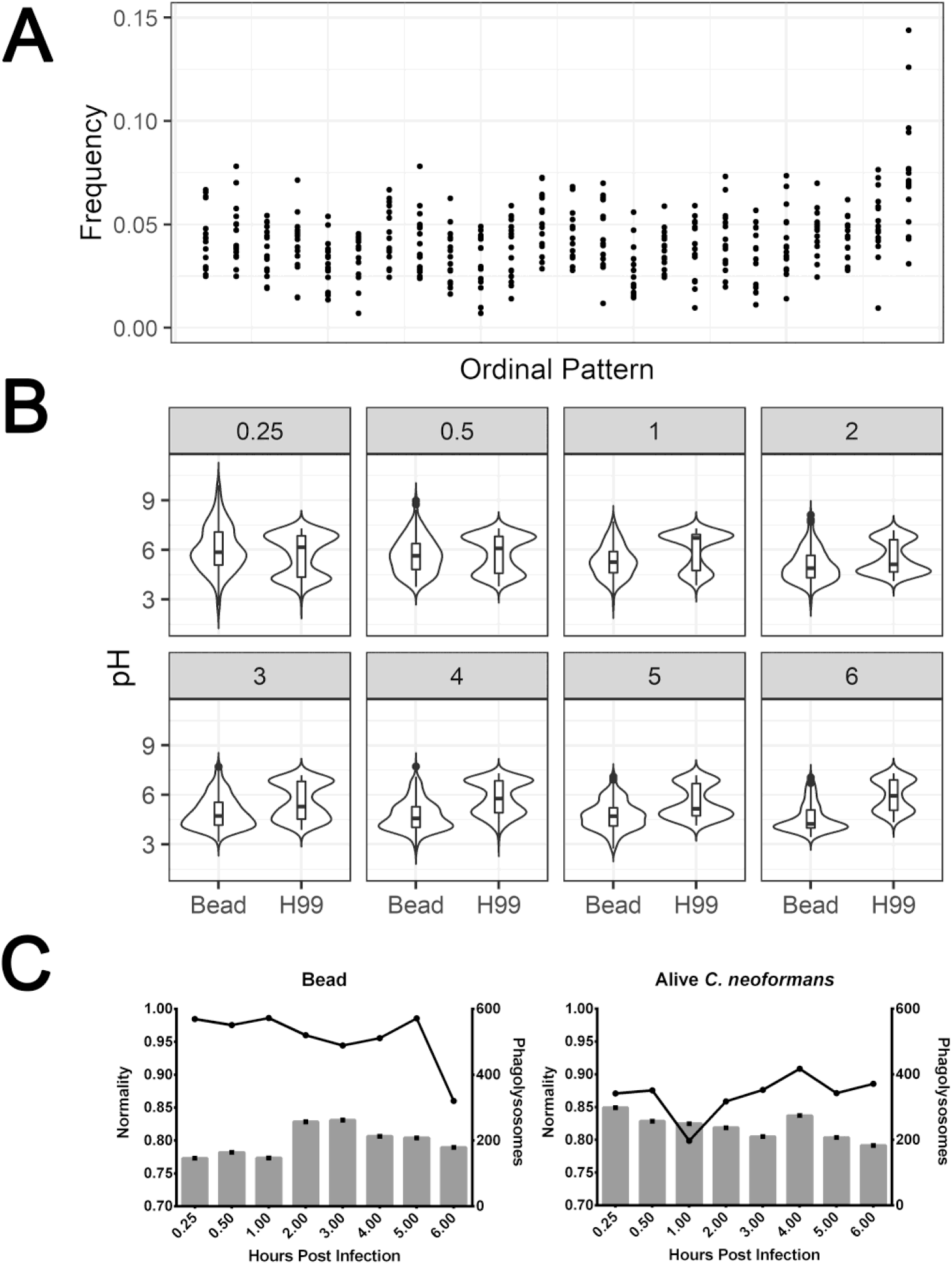
Isolated human monocytes acidify stochastically and approximate normality. Human macrophages were infected with inert beads or live *C. neoformans*, and their pH analyzed at various timepoints. **A.** Ordinal Pattern analysis for all conditions. All patterns exist at non-zero frequencies for all timepoints. **B.** Distributions of phagolysosome pH at different timepoints. **C.** Shapiro-Wilk normality and total sample count for each condition.

### *Pathogens* skew phagolysosome acidification towards conditions less favorable to the macrophage

Cryptococcal cells buffer the phagolysosome pH toward 5.5, a value optimal for yeast growth^14^. We therefore hypothesized that *C. neoformans* could use this buffering capability to disrupt the host acidification strategy. We found that phagolysosomes containing live *C. neoformans* and live *C. gattii* acidified to distributions of pH less normally distributed and more permissive to general fungal replication by increasing the average observed phagolysosomal pH.

To probe host fitness with regard to *C. neoformans* specifically, we modified our model by declaring the inhibitory pH as ≤ 4, which inhibits *C. neoformans* replication^34^, and calculated the likelihood of each population of macrophages to achieve an inhibitory pH, assuming phagolysosome pHs were normally distributed around the observed mean and standard deviation. Here we analyzed the combined data for all time intervals, reasoning that in actual infection interactions between *C. neoformans* and host macrophages early phagosomes would be important as well, rather than focusing only on mature phagolysosomes.

We estimated the proportions of inhibitory phagolysosomes and found that bead-containing phagolysosomes were more likely to acidify to ≤ 4, followed by heat-killed and *Δure1 C. neoformans*, followed by live *C. neoformans*, with capsule deficient *Cap59* being particularly likely, and trained macrophages or *C. gattii* containing phagolysosomes being particularly unlikely to achieve inhibitory pH (Fig 9A). Our actual data was not normally distributed with non-bead samples though, as *C. neoformans* actively modulates phagolysosomal pH. The starkest difference we observed is that, in reality, killed and live *C. neoformans* phagolysosomes had the same proportion of inhibitory phagolysosomes, suggesting the capsule has a more significant effect on pH modulation than we initially expected. This finding is corroborated by the high proportion of inhibitory phagolysosomes in *Cap59* containing phagolysosomes, a strain incapable of modulating pH since it has no capsule (Fig 9C). Additionally, the expected and observed increased likelihood of bead containing phagolysosomes to inhibit *C. neoformans* replication compared to live *C. neoformans* phagolysosomes was consistent between murine and human cells (Fig 9B,D).

**Figure 9.**
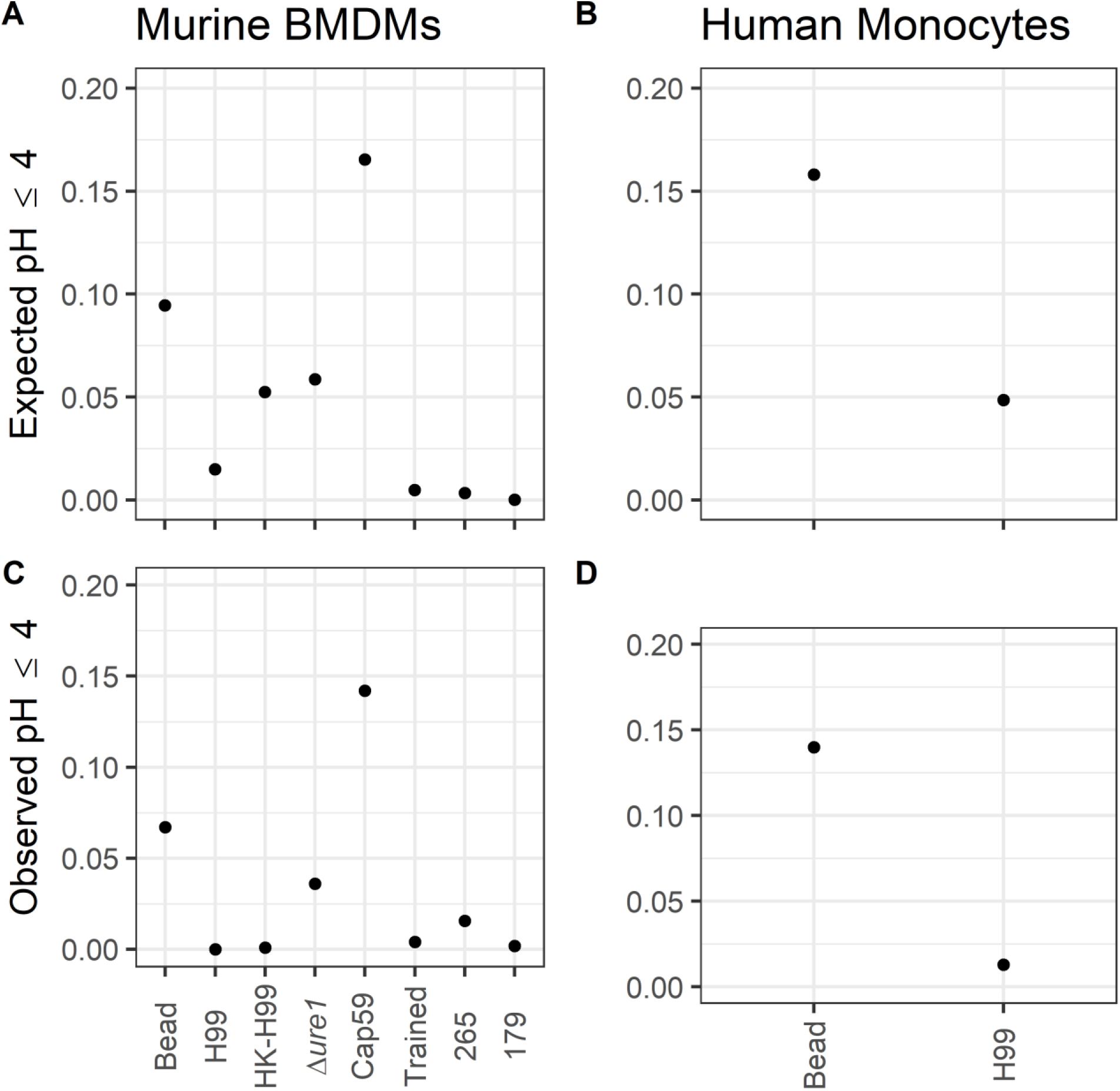
Expected and observed likelihood of phagolysosomes to achieve pH ≤ 4. **A.** The expected proportions of murine BMDM phagolysosomes to achieve pH ≤ 4 assuming a normal distribution based on all observed data. **B.** The expected proportions of human macrophage phagolysosomes to achieve pH ≤ 4 assuming a normal distribution based on the observed data. Expected proportions of macrophages to achieve pH ≤ 4 were calculated by assuming a normal distribution centered around a mean and standard deviation calculated from the observed data. **C.** The observed proportions of murine BMDM phagolysosomes to achieve pH ≤ 4. **D.** The observed proportions of human macrophage phagolysosomes to achieve pH ≤ 4.

To probe whether this phenomenon was applicable to pathogens other than *C. neoformans,* we repeated this analysis with published phagolysosomal pH data from *Mycobacterium avium*^35^. We found that like *C. neoformans,* live *M. avium* bacteria modified their resident phagolysosomal pH to be more favorable toward them. In contrast, the estimated mean log fitness for the host macrophage population increased when they ingested killed mycobacteria, which are unable to modulate pH (SFig 11).

## Discussion

The complexity and sequential nature of the phagolysosomal maturation process combined with the potential for variability at each of the maturation steps, and the noisy nature of the signaling networks that regulate this process, have led to the proposal that each phagolysosome is a unique and individual unit^36^. In fact, the action of kinesin and dynein motors that move phagosomes along microtubules exhibits stochastic behavior, adding an additional source of randomness to the process^37^. Hence, even when the ingested particle is a latex bead taken through one specific type of phagocytic receptor there is considerable heterogeneity in phagolysosome composition, even within a single cell^36^. Since the phagolysosome is a killing machine used to control ingested microbes, this heterogeneity implies there will be differences in the microbicidal efficacy of individual phagolysosomes. This variability raises fundamental questions about the nature of the dynamical system embodied in the process of phagosomal maturation.

In this study, we analyzed the dynamics of phagolysosome pH variability after synchronized ingestion of live yeast cells, dead yeast cells, and latex particles. We sought to characterize the acidification dynamics as either stochastic—an inherently unpredictable process with identical starting conditions yielding different trajectories in time, or deterministic—a theoretically predictable process with identical starting conditions leading to identical trajectories. In particular, we focused our analysis on differentiating stochastic vs. chaotic signatures in the trajectories of phagolysosomal pH. While both dynamics might yield highly divergent trajectories for similar starting conditions (i.e. only one of 100 variables differing by only a minuscule amount), a chaotic system is inherently deterministic, such that if strictly identical starting conditions were replicated, the same trajectory would follow from those conditions each time. A chaotic system is defined as one so sensitive to initial conditions that, in practice, initial conditions cannot be replicated precisely enough to see these same trajectories followed. The dynamical signatures of such systems are unique and can be differentiated from that of other deterministic or stochastic dynamics.

Irrespective of the nature of the ingested particle, we observed that the distribution of the increment of phagolysosomal pH reduction was random, indicative of a stochastic process. We found no evidence that phagosome acidification was a chaotic process. Systems in which a large number of variables each contribute to an outcome tend to exhibit ‘noise’, which gives them the characteristics of a stochastic dynamical system. Additionally, while particle size and shape affects ingestion time^38^ the mean time to ingestion for beads and *C. neoformans* particles have been shown to be 2.5 and 4.18 min, respectively^38,39^.

Since phagocytosis was synchronized and our observations were made over a period of hours, it is unlikely that the effects described here are due to noise from differences in uptake time differences. In this regard, our finding that phagolysosomal pH demonstrates stochastic features is consistent with our current understanding of the mechanisms involved. Quantile-Quantile (Q-Q) plots revealed that most phagolysosomal pH distributions in this study manifested significant deviations from normality in several instances. Specifically, this stochastic normal distribution was generated at the phagolysosomal level, as evident by the fact that different phagosomes within the same macrophage manifested different pH values. Hence, each macrophage contains phagolysosomes with a different pH rather than each macrophage containing multiple of the same pH, such that the normal distribution observed was generated at the organelle level. The most normally distributed pH sets were those resulting from the ingestion of latex beads, particles that cannot modify the acidity of the phagolysosome. We note that for the three *C. gattii* strains the pH distributions revealed more skewing in Q-Q plots than for the H99 *C. neoformans* strain. Although the cause of this variation is not understood and the strain sample size is too small to draw firm conclusions, we note that such variation could reflect more microbial-mediated modification of the phagolysosomal pH by the *C. gattii* strains. In this regard, the capsular polysaccharide of *C. gattii* strains has polysaccharide triads that are more complex^40^ and, given that the cryptococcal polysaccharide capsule contains glucuronic acids that can modify phagolysosomal pH via acid-base properties^17^, it is possible that this skewing reflects differences in phagolysosome to phagosome capsular effects.

We attempted to separate the dynamics of phagolysosomal maturation from acidification by investigating the accumulation of phagolysosomal maturation markers, EEA1 and V-ATPase after ingestion. Analyzing the same time intervals where pH populations stabilized, namely 1 to 4 h post ingestion, we found no forbidden ordinal patterns, suggesting that phagosome maturation was also a stochastic process. However, the acquisition of these two maturation markers did not approximate a normal distribution at any of the four time intervals, with all samples manifesting heavy right skewing. This is especially interesting considering V-ATPase is responsible for pumping protons into the phagolysosome and maintaining acidity. If the number of V-ATPase molecules translated directly to the number of protons pumped into the phagolysosome we would expect correlation between V-ATPase staining intensity and pH, and thus expect normal distributions in both. The different dynamics observed with V-ATPase immunofluorescence implies the pH heterogeneity is regulated by additional mechanisms. For example, it is possible that the efficacy of the V-ATPase pumps on the phagolysosomal membrane differs from pump to pump and that these differences also contribute to the distribution of phagolysosomal pHs observed.

For most microbes, maintenance of an acidic environment in the phagolysosome is critically determinant on the integrity of the phagolysosomal membrane, keeping protons in the phagolysosomal lumen whilst excluding more alkaline cytoplasmic contents. For example, with *C. albicans,* the rupture of the phagolysosomal membrane is followed by rapid alkalization of the phagolysosomal lumen^41^. For *C. neoformans*, phagolysosomal integrity is compromised by secretion of phospholipases that damage membranes as well as the physical stress on membranes resulting from capsular enlargement in the phagolysosome^14^. However, for *C. neoformans*, loss of phagolysosomal membrane integrity does not immediately result in loss of phagolysosomal acidity, which is attributed to buffering by glucuronic acid residues in the *C. neoformans* capsule^17^. Adding to the complexity of the *C. neoformans*-macrophage interaction is the fact that the phagolysosomal pH in the vicinity of 5.5 matches the optimal replication pH for this fungus^15^, which can be expected to place additional stress on the organelle membrane through the increased volume resulting from budding cells. Treating macrophages with chloroquine, which increases phagosomal pH^42^, potentiates macrophage antifungal activity against *C. neoformans*^43^. Hence, phagosomal acidification does not inhibit *C. neoformans* replication but is critical for activation of mechanisms involved in antigen presentation^44^. In the cryptococcal-containing phagolysosome the luminal pH is also likely to reflect a variety of microbial-mediated variables which include ammonia generation from urease, capsular composition, and the integrity of the phagolysosomal membrane.

Analysis of the normality of phagolysosomal pH distributions as a function of time by the Shapiro-Wilk test produced additional insights into the dynamics of these systems. Phagolysosomes containing inert beads manifested pH distributions that met criteria for normality at most time intervals after 1 h post infection (HPI). We hypothesize that at 0.25 and 0.5 HPI there are two populations of phagolysosomes, mature and maturing, with the latter having not yet fully acidified and thus resulting in the observed bimodal distributions. Additionally, it has been shown that phagolysosomes of macrophages undergo active alkalization, regulated in part by NOX2 activity^45–47^. It is likely that these early time points veer away from normality due to a combination of phagolysosomes maturing at a different rate and a subpopulation of phagolysosomes which are actively alkalized, both contributing noise to early phagolysosomal dynamics. In contrast, the pH distribution of phagolysosomes containing dead *C. neoformans* cells initially veered away from normality at 15 min, 30 min, and 1 h but in later time intervals approached normality, meeting the criteria for normality at 3 h. One interpretation of this result is that the process of phagocytosis is itself a randomizing system with Gaussian noise resulting from phagolysosome formation and kinetics of the initial acid-base reactions between increasing proton flux and quenching glucuronic acids in the capsular polysaccharide. With time, the titration is completed as all glucuronic acid residues are protonated, as dead cells did not synthesize additional polysaccharide, which moved the phagolysosomal pH distribution towards normality. A similar effect may have occurred with strains R265, W179, and *ure1Δ*. Convergence to or away from normality could reflect a myriad of such variables affecting phagolysosomal pH including the intensity of acidification, the volume of the phagolysosome (largely determined by the yeast capsule radius), the glucuronic acid composition of the capsule, the production of ammonia by urease, and the leakiness of the phagolysosome to cytoplasmic contents with higher pH. Although our experiments cannot sort out the individual contributions of these factors, they suggest that, in combination, they produce Gaussian noise effects that push or pull the resulting distribution to or from normality. Additionally, human phagolysosome acidification dynamics resembled those of mouse cells but noted significant differences in the distributions of phagolysosomal pH between individual human donors. This donor-to-donor variation could reflect differences in polymorphisms in Fc receptor genes or other genetic variables and is an interesting subject for future studies.

When a phagocytic cell ingests a microbe, it has no information as to the pH range tolerated by the internalized microbe. A stochastic dynamical process for phagolysosomal acidification could provide phagocytic immune cells and their hosts with the best chance for controlling ingested microbes. The acidic pH in the phagolysosome activates microbicidal mechanisms and acidity is not generally considered a major antimicrobial mechanism in itself. However, our analysis of pH tolerances of 27 pathogenic microbes revealed that the majority are inhibited by phagolysosomal pH with the caveat that some, like *neoformans*^18^ and *Salmonella typhimrium*^48^, thrive in acidified phagolysosomes. On the other hand, a less acidic phagosomal pH is conducive to intracellular survival for *M. tuberculosis*^49^. During an infectious process when the immune system confronts numerous microbial cells the random nature of the final phagosomal pH will result in some fraction of the infecting inoculum being controlled, and possibly killed, by initial ingestion allowing antigen presentation. In this regard, the mean number of bacteria in the phagolysosomes and cytoplasm of macrophages infected with the intracellular pathogen *Franciscella tularensis* exhibits stochastic dynamics^50^, which in turn could result from the type of stochastic processes in phagolysosome formation noted here. Hence, chance in phagolysosomal pH acidification provides phagocytic cells with a mechanism to hedge their bets such that the stochastic nature of the process is itself a host defense mechanism.

In biology, bet-hedging was described by Darwin as a strategy to overcome an unpredictable environment^51^, which is now known as diversified bet-hedging: diversifying offspring genotype to ensure survival of at least some individuals at the expense of reducing the mean inclusive fitness of the parent. The main idea behind any bet-hedging strategy, under the assumption of multiplicative fitness, is that to maximize long-term fitness, an organism must lower its variance in fitness between generations^52–54^. For example, varying egg size and number in a clutch can bank against years with a hostile environment in a form of diversified bet-hedging^55^. During any given “good year” fewer of the offspring will thrive because some are specifically designated for survival in “bad years”, drastically increasing fitness during bad years at the cost of a slight fitness reduction during good. Our observations suggest that, as a population, macrophages perform a bet-hedging strategy by introducing a pH level as inhospitable to pathogens as possible, while still maintaining biologically possible levels. However, such an approach could select for acid-resistant microbes. Our observation suggests that to avoid an arms-race, the macrophage not only lowers the pH level to a level that is unfavorable to most microbes, since the reduction in pH level reduces the optimal growth condition to around 70% of the optimum (see Figure 3 red line under the blue shaded curve), but it also introduces randomness in the achieved pH, such that ingested microbes are less likely to adapt to the potentially hostile environment. In other words, tightly controlled pH reduction by the host without increased variation, might introduce an evolutionary arm-race between pathogens and their host cells, leading towards deleterious outcome of selecting microbial acid resistance. A similar evolutionary arms race would occur between environmental *C. neoformans* and the amoebas who prey upon them. Previous studies indicate that acidification in amoebas closely resembles that of macrophages with similar final pH and time to acidification^56,57^. Thus, phagocytic predators in soils would face the same problem as macrophages in not knowing the pH tolerance of their prey and it is conceivable that they employ a similar defense strategy to that observed in macrophages, likely to have been honed in by eons of selection in soil predator-prey interactions.

Additionally, our model shows that even at the lower extreme mean pH, macrophage populations still benefit in the long run by increasing their phagolysosomal pH variance. We note that the pH of other mammalian fluids such as that of the blood are tightly regulated such that their physiological variance is very small^58^. For example, human plasma pH averages 7.4 with a range of only 0.05 units. Hence, organisms can maintain tight pH control when it is physiologically important, implying that the comparatively large range of phagolysosomal pHs measured in all conditions studied is a designed feature of this system. In other words, one can envision the macrophage’s first line of defense of bet-hedging phagolysosomal pH as a roulette wheel, where the payout is likelihood of macrophage survival and the placed bet is a range of possible pH. Placing multiple bets across the table (increasing standard deviation of pH distribution) increases the chance of winning at the cost of a lower payout (mean fitness), resulting in a more profitable long-term strategy (increased mean log fitness). In fact, the most profitable roulette strategy is broad color bets with lower payouts but the best winning probability. The most profitable betting strategy would of course be to play Blackjack instead, but macrophages do not have that luxury.

We observed that this bet hedging strategy was displayed in M1 polarized macrophages, dependent on a normal distribution of phagolysosomal pH with a high variance. M2 macrophages, which acidify with different dynamics to M1^59^, do not acidify to a normal distribution and thus do not engage in this bet hedging strategy. Additionally, we found that M2 skewed macrophage populations on average acidify to a higher pH than M1 skewed populations and are overall more favorable to *C. neoformans*, an observation supported by previous literature^59,60^. This hypothesis, while requiring more investigation, may help explain why M2 skewed macrophages are unable to control *C. neoformans* infection, as their phagolysosomes acidify to a pH range that is optimal for *C. neoformans* growth^15^.

Additionally, intracellular pathogens have developed their own ways to disrupt or game the macrophage betting system. For *C. neoformans*, which can manipulate phagolysosomal pH through several mechanisms, our model shows that the fungal-mediated changes in the distribution of phagolysosomal pH favors the pathogen overall, disrupting the bet hedging system and resulting in lower macrophage population fitness. This phenomenon can also be observed with *M. avium* in which previously reported phagolysosomal pH values^35^ show a marked increase in mean log fitness when the ingested pathogen is killed and unable to modulate pH. This finding is emphasized by our analysis of phagolysosomes whose pH has been pharmacologically manipulated to a region favorable to pathogens. We found that disrupting the macrophage betting system this way led to a drastically reduced overall mean log fitness of the macrophage population.

Given that acid has potent antimicrobial properties, one might wonder why phagolysosomal acidification does not reach even lower and more acidic pHs. There are several explanations for observed lower limits in pH. Acidification is achieved by pumping protons into the vacuole and achieving lower pHs against an ever-increasing acidity gradient could prove thermodynamically difficult. There is also evidence that the integrity of the cell membrane lipid bilayer is compromised by acidity at pHs of 3 and below^61,62^ which could promote leakage of phagolysosomal contents into the cytoplasm with damage to the host cell. Consequently, we propose that a larger variation in macrophage phagolysosomal pH acts as a diversified bet-hedging strategy against the stochasticity of the potential pH tolerances of ingested microbes within the physiological limits of achievable acidification. This hypothesis is supported by simulated data in which an increase in standard deviation of the pH distribution slightly lowers the expected mean but significantly decreases the standard deviation of host survival. Fully analyzing the consequences and evolutionary tradeoff of this strategy would require a closer analysis that considers the costs and benefits with regard to the host. Though outside the scope of our current work, our observations suggest this line of investigation for future studies.

Our results delineate new avenues for investigation. Most perplexing is how a random phagolysosomal pH is established and maintained. One can imagine various mechanisms including variation in the number of ATPase molecules and/or differences in activity of individual pumps depending on location and adjoining structures. To resolve this would require sophisticated technology allowing the measurement of pH in individual phagolysosomes as a function of pump occupancy and efficacy. Such techniques are likely beyond the current technological horizon but suggests new fertile areas of scientific investigation. From a clinical perspective, a drug that increases variation in phagolysosomal pH could be useful in enhancing macrophage anti-microbial efficacy in situations that one cannot anticipate which specific microbes will be encountered. On the other hand, drugs that specifically modulate pH may be useful in situations with specific microbes that express marked acid/base tolerance. In this regard, chloroquine alkalizes *C. neoformans* phagolysosomes, moving away from optimal inhibitory pH. This drug has been shown to enhance macrophage activity against *C. neoformans*^9,63^, an outcome predicted by our model.

In summary, we document that phagolysosomal acidification, a critical process for phagocytic cell efficacy in controlling ingested microbial cells, manifests stochastic dynamics that permit a bet-hedging strategy for phagocytic cells ingesting microbes of unknown pH tolerance. These observations establish that the use of bet-hedging strategies in biology extends to the sub-organism level to involve cells and their organelles. This, in turn, implies a significant role for chance in the resolution of conflict between microbes and host phagocytic cells in individual phagolysosomes. We have recently argued that chance is also a major determinant of individual susceptibility to infectious diseases at the organismal level^64^. Variability in the outcome of infectious disease between individual hosts may reflect the sum of innumerable chance events for host-microbe interactions at the cellular level, which include the process of phagosome acidification.

## Methods

### Cell Strains and Culture Conditions

*Cryptococcus gattii* species complex strains R265, WM179, and WM161 were obtained from ATCC (Manassas, VA), and *Cryptococcus neoformans* species complex serotype A strain H99 as well as *ure1Δ* (lacking urease, derived from H99) were originally obtained from John Perfect (Durham, NC). The strains were stored at 80°C.Frozen stocks were streaked onto Sabouraud dextrose agar (SAB) and incubated at 30°C. Liquid suspensions of cryptococcal cultures were grown in SAB overnight at 30°C. Cryptococcal cultures were heat killed by incubating at 65 °C for 1 h.

Macrophage cells were either mouse bone marrow-derived macrophages (BMDM) obtained from 6 week old C57BL/6 female mice from The Jackson Laboratory (Bar Harbor, ME), J774.16 macrophage-like cells, or human monocytes isolated from PMBCs. BMDMs were isolated from hind leg bones and for differentiation were seeded in 10 cm tissue culture treated dishes (Corning, NY) in Dulbecco’s Modified Eagle Medium (DMEM, Corning) with 10% FBS (Atlanta Biologicals, Flowery Branch, GA), 1% nonessential amino acids (Cellgro, Manassas, VA), 1% penicillin-streptomycin (Corning), 2 mM Glutamax (Gibco, Gaithersburg, MD), 1% HEPES buffer (Corning), 20% L-929 cell conditioned supernatant, 0.1% beta-mercaptoethanol (Gibco) for 6 days at 37 °C and 9.5% CO_2_. BMDMs were used for experiments within 5 days after differentiation. J774.16 cells were cultured in DMEM with 10% FBS, 1% nonessential amino acids, 10% NCTC109 (Gibco), and 1% penicillin-streptomycin at 37 °C with 9.5% CO_2_. For human peripheral blood mononuclear cells (hPBMCs), CD14+ monocytes were isolated using Dynabeads Untouched Human Monocytes Kit (Thermo Fisher Scientific) according to the manufacturer protocol. The isolated cells were differentiated in RPMI-1640 medium (RPMI) with 10% FBS (Atlanta Biologicals, Flowery Branch, GA), 1% penicillin-streptomycin (Corning), and 25 ng/mL of human granulocyte-macrophage colony-stimulating factor (GM-CSF, Sigma-Aldrich) for five days. Human cells were further cultured in RPMI (RPMI) with 10% FBS (Atlanta Biologicals, Flowery Branch, GA), 1% penicillin-streptomycin (Corning), and activated with 0.5 ug/mL lipopolysaccharide (LPS; Sigma-Aldrich) and 10 ng/mL interferon gamma (IFN-γ; Roche) for M1 polarization or 20 ng/mL IL-4 for M2 polarization.

### Phagolysosomal pH measurement

Phagolysosomal pH was measured using ratiometric fluorescence imaging involving the use of pH-sensitive probe Oregon green 488 as described in prior studies^15^. The pH values analyzed here were collected in part during prior studies of *C. neoformans*-macrophage interactions^14–16^. Briefly, Oregon green 488 was first conjugated to monoclonal antibody (mAb) 18B7, which binds *C. neoformans* capsular polysaccharide, using Oregon Green 488 Protein Labeling Kit (Molecular Probes, Eugene, OR). The labeling procedure was done by following the manufacture’s instruction. BMDMs were plated at a density of 1.25 × 10^5^ cells/well or differentiated human macrophages were plated at a density of 2.5 × 10^5^ cells/well on 24-well plate with 12 mm circular coverslip. Cells were activated with 0.5 μg/ml LPS and 100 U/ml IFN-γ or 20 ng/mL IL-4 as previously described at 37 °C in a 9.5% CO_2_ (BMDM) or 5% CO_2_ (Human macrophage) atmosphere overnight. Prior to infection, 2 d old live, heat killed H99, R265, WM179, ure1, cap59, or anti-mouse IgG coated polystyrene bead (3.75 × 10^6^ cells or beads/ml) were incubated with 10 μg/ml Oregon green conjugated mAb 18B7 for 15 min. Macrophages were then incubated with Oregon green conjugated mAb 18B7-opsonized particles in 3.75 × 10^5^ cryptococcal cells or beads per well (BMDM) or 2.5 × 10^5^ cell/well (Human macrophage). For drug treatment experiments the macrophage cell media was supplemented with 3 μM chloroquine. Cells were either centrifuged immediately at 350 x g for 1 min or incubated at 4 °C for 30 min to synchronize ingestion and cultures were incubated at 37 °C for 10 min to allow phagocytosis. Extracellular cryptococcal cells or beads were removed by washing three times with fresh medium, a step that prevents the occurrence of new phagocytic events. As an additional safeguard against new phagocytic events fresh media was supplemented with AlexaFluor 568 conjugated mAb 18B7 for 1 h to label extracellular particles. Samples on coverslip were collected at their respective time points after phagocytosis by washing twice with pre-warmed HBSS and placing upside down on MatTek petri dish (MatTek, Ashland, MA) with HBSS in the microwell. Images were taken by using Olympus AX70 microscopy (Olympus, Center Valley, PA) with objective 40x at dual excitation 440 nm and 488 nm, and emission 520 nm. Images were analyzed using MetaFluor Fluorescence Ratio Imaging Software (Molecular Devices, Downingtown, PA). Fluorescence intensities were used to determine the ratios of Ex488 nm/Ex440 nm that were converted to absolute pH values using a standard curve where the images are taken as above but intracellular pH of macrophages was equilibrated by adding 10 μM nigericin in pH buffer (140 mM KCl, 1 mM MgCl_2_, 1 mM CaCl_2_, 5 mM glucose, and appropriate buffer ≤ pH 5.0: acetate-acetic acid; pH 5.5-6.5: MES; ≥ pH 7.0: HEPES. Desired pH values were adjusted using either 1 M KOH or 1 M HCl). The pH of buffers was adjusted at 3-7 using 0.5-pH unit increments.

### Immunofluorescence Microscopy

BMDMs were seeded on 12 mm circular coverslips in 24 well tissue culture plates at 2.5 * 10^5^ cells per well. Cells were activated with 0.5 μg/ml lipopolysaccharide and 100 U/ml interferon gamma at 37 °C in a 9.5% CO_2_ atmosphere overnight. Anti-mouse IgG coated beads were opsonized with 10 μg/mL and added to cells at MOI 1. Phagocytosis was synchronized by centrifuging the plate at 350 x g for 1 min before being incubated at 37 °C. At each timepoint media was replaced with 4% paraformaldehyde (PFA) and incubated for 10 min at room temperature to fix cells. PFA was removed and cells were washed three times with 1 mL PBS (1X). Coverslips were blocked for 1 h at room temperature with 2% BSA in PBS. Primary incubation was performed with Rb αEEA1 (ThermoFisher Scientific MA5-14794) at 1:100 dilution or Rb αV-ATPase (ThermoFisher Scientific PA5-29899) at 1:100 dilution in blocking buffer for 1 h at room temperature. Coverslips were washed with blocking buffer before secondary incubation with Goat αRb AlexaFluor 488 (1:100) in blocking buffer for 1 h at room temperature. Coverslips were washed once more with dH_2_O and mounted on glass slides using Prolong Gold mounting agent. Slides were imaged via Zeiss Axiovert 200M microscope and intensity was analyzed using Zeiss Lite Blue Version. A region of interest was generated by outlining the ingested bead via phase contrast channel then measuring mean fluorescence intensity of the secondary antibody.

### Trained macrophage experiments

BMDMs were plated at a density of 2 × 10^5^ cells/well on 24-well plate with 12 mm circular coverslip. Cells were activated with 0.5 μg/ml LPS and 100 U/ml IFN-γ; Roche and incubated at 37 °C in a 9.5% CO_2_ atmosphere overnight. Prior to the infection, H99 were stained with 0.01% Uvitex 2B for 10 min. BMDMs were then infected with H99 (2 × 10^5^ cells/well) in the presence of 10 μg/ml 18B7. After 1 h of infection, 5 μg/ml of amphotericin B were added to each well and the culture were incubated for overnight. On the following day, the cultures were washed three times with PBS, and incubated in PBS for 2 h. After the incubation, the cultures were further washed three times with PBS. A fresh overnight culture of H99 were incubated with 10 μg/ml Oregon green conjugated mAb 18B7 for 15 min. Macrophages were then incubated with Oregon green conjugated mAb 18B7-opsonized H99 in 2 × 10^5^ cells per well. Cells were centrifuged immediately at 350 x g for 1 min to synchronize ingestion. Phagolysosomal pH were then measured using Olympus AX70 microscopy.

### Time-lapse imaging and intracellular replication

The time of intracellular replication here were collected in time-lapse imaging during prior studies of *C. neoformans*-macrophage interactions^15^. For imaging BMDM (5 × 10^4^ cells/well) were plated on poly-D-lysine coated coverslip bottom MatTek petri dishes with 14mm microwell (MatTek). Cells were cultured in completed DMEM medium and stimulated with 0.5 μg/ml LPS and 100 U/ml IFN-γ overnight at 37 °C with 9.5 % CO_2_. On the following day, macrophages were infected with cryptococcal cells (H99 or ure1; 1.5 × 10^5^ cells/well) opsonized with 18B7 (10 μg/ml). After 2 h incubation to allow phagocytosis, extracellular cryptococcal cells were removed by washing the culture five times with fresh medium. Images were taken every 4 min for 24 h using a Zeiss Axiovert 200M inverted microscope with a 10x phase objective in an enclosed chamber at 9.5 % CO_2_ and 37 °C. The time intervals to initial replication of individual cryptococcal cells inside macrophage were measured in time-lapse imaging.

### Data Processing

Phagolysosome pH intervals were calculated by subtracting measured pH levels of phagolysosomes from a starting pH of 7.2, the pH of the surrounding media, and individual interval measurements were concatenated into a single dataset for each time point examined.

### Data analysis

Discrimination of deterministic vs. stochastic dynamics was achieved using the previously characterized permutation spectrum test^21^. In this method, the processed datasets were segmented into overlapping subsets of 4 data points using a sliding window approach, as detailed in figure 4, and assigned 1 of 24 (4!) possible ordinal patterns based on the ordering of the 4 terms in the subset. The frequencies with which each unique ordinal pattern occurred in the dataset were then calculated and plotted. Deterministic dynamics were characterized by the occurrence of “forbidden ordinals”, equal to ordinal patterns that doesn’t occur in the dataset whereas stochastic dynamics were characterized by the presence of all ordinals. Measured phagolysosomal pHs were subtracted from an initial pH value (7.2) based on cell media pH and placed in a vector. Subsets of 4 data points were generated using a sliding window approach in which the first four values were grouped, the window shifted by one, and the subsequent set of 4 values grouped. Each subset was prescribed an “ordinal pattern” based on the relative values of the data points in the subset to each other with, for instance, the lowest value assigned a “0” in the ordinal pattern and the highest a “3”. Further characterization of deterministic dynamics was achieved using the previously characterized point count plot^65^, in which periodic vs. chaotic dynamics were differentiated based on the distribution of “peaks” in the calculated power spectrum of each dataset. Power spectrums were estimated with Matlab’s Lomb-Scargle power spectral density (PSD) estimate function and subsequently normalized. From the normalized power spectrum, “point count plots” were generated by counting the number of peaks above a set threshold—the point threshold—with values of the point threshold ranging from 0 to 1. Periodic dynamics were characterized by “staircase” point count plots whereas chaotic dynamics were characterized by point count plots with a decreasing exponential shape.

### Distribution and normality analysis

Each set of sample data was fit to a series of distributions using the R package “fitdistrplus” with default parameters for each distribution type, generating the histograms and Quantile-Quantile (Q-Q) plots. Normality and significance were calculated via the base R Shapiro-Wilk test.

### Generating and Analyzing Simulations

Let f(x) be the probability distribution of the host’s pH. For simplicity, we assume that f is Normal(μ, σ). Let F(x) be the corresponding cdf. At every time-point, we randomly sample a pathogen i from a set of pathogens. The pathogen has a range (*a*_*i*_,*b*_*i*_) of viable pH, which differs among pathogens. Minimum tolerable pH values were obtained from the literature as described, while a constant value of 8 was used for maximum since most pathogens were inhibited as high pH as well as phagolysosomes being unlikely to alkalize. The host, in turn, randomly chooses a pH by sampling from the distribution f(x). If the host’s pH x falls in the interval (*a*_*i*_,*b*_*i*_), then there is a probability p of the host dying. We can think of the host’s survival rate ρ as its fitness, which is a random variable that depends on the randomness in pathogens. As a function of i, we have ρ(i)=p*(F(*b*_*i*_)-F(*a*_*i*_)); the random variable ρ assumes the values ρ(i) with equal probability. Additionally, since biologically relevant phagolysosome pH values will only fall within a certain range, we decided to limit the possible pH values between 2 and 7.2 (pH of associated cell media). Any values generated outside those limits were instead recorded as their respective limit. Thus, distributions become less normally distributed at the extremes, sacrificing normality for biological practicality, as real phagolysosomes would not alkalize or be likely to acidify below 2. In these situations, we calculated outputs based on 1,000 replicates of 10,000 simulated phagolysosomes each.

## Author Contributions

Conceptualization, Q.D., K.S., A.V, A.C; Methodology: Q.D., K.S., Y.S., M.F., Y.S., C.Y., A.B., A.C.; Software, Q.D., K.S., Y.S., A.V.; Formal Analysis, Q.D.; Investigation, Q.D., K.S., Y.S., M.F., Y.S., C.Y., I.Y., C.M.DLR, J.F., A.B., A.C.; Writing – Original Draft, Q.D., K.S., M.F, A.B., A.C.; Writing – Review & Editing, Q.D., K.S., Y.S., M.F., Y.S., C.Y., I.Y., C.M.DLR, J.F., A.B., A.C.; Visualization, Q.D.

## Acknowledgements

Arturo Casadevall was supported by grants R01HL059842, 5R01AI033774, 5R37AI033142, and 5R01AI052733. Quigly Dragotakes is supported by a fellowship from the Achievement Rewards for College Scientists (ARCS) Foundation Metro-Washington Chapter as well as a National Institutes of Health T32.

---

**Supplemental Figure 1.**
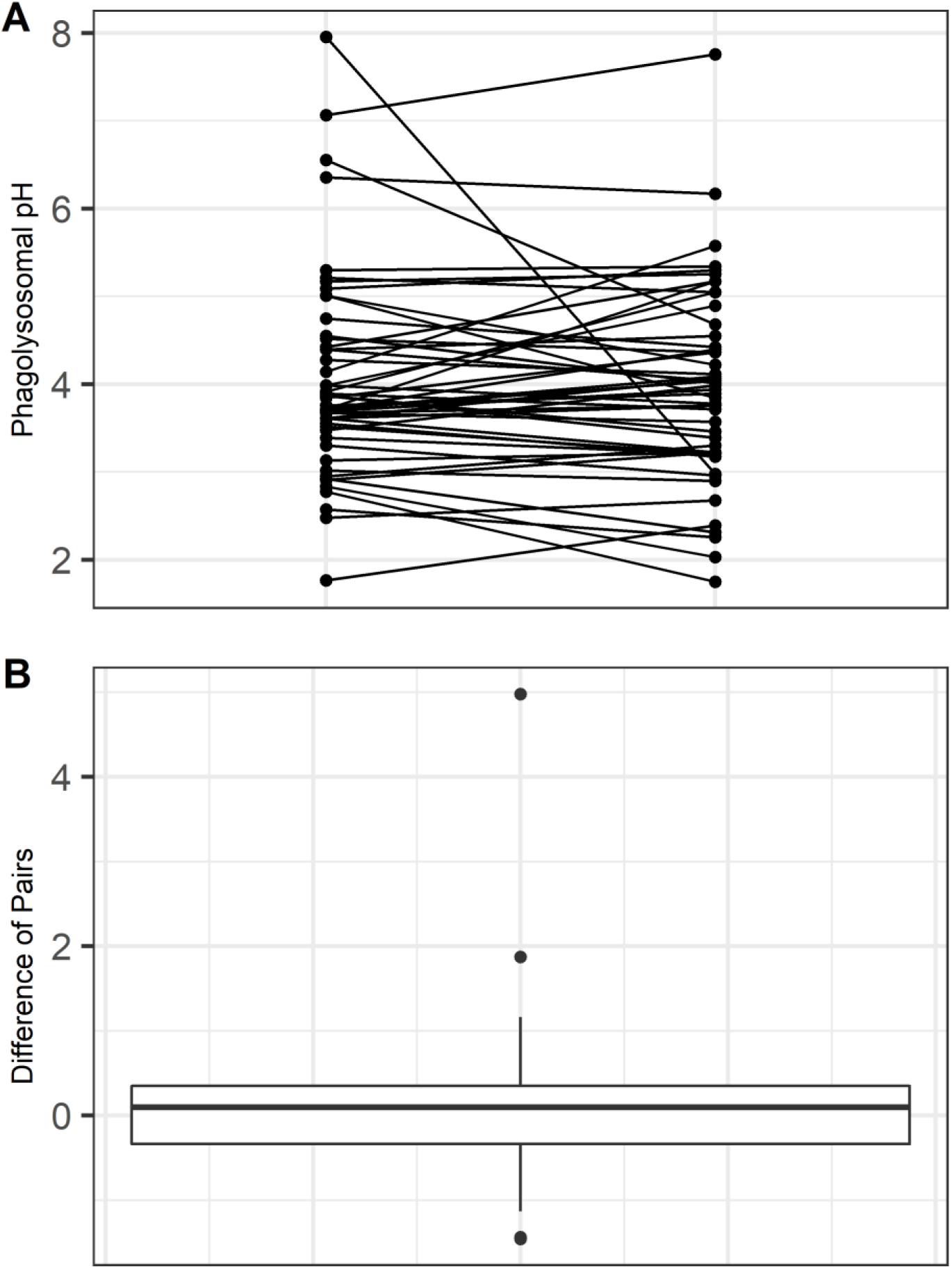
Analysis of 80 pairs of phagolysosomes within single cells. **A.** Measured phagolysosomal pH of multiple beads located in the same macrophage but non-proximal phagolysosomes. Lines indicate particles within the same macrophage. Phagolysosomes were measured in no particular order. **B.** Differences of each phagolysosomal pH pair approximate a normal distribution centered around 0 with non-zero values despite being generated within the same macrophage at a synchronized starting time.

**Supplemental Figure 2.**
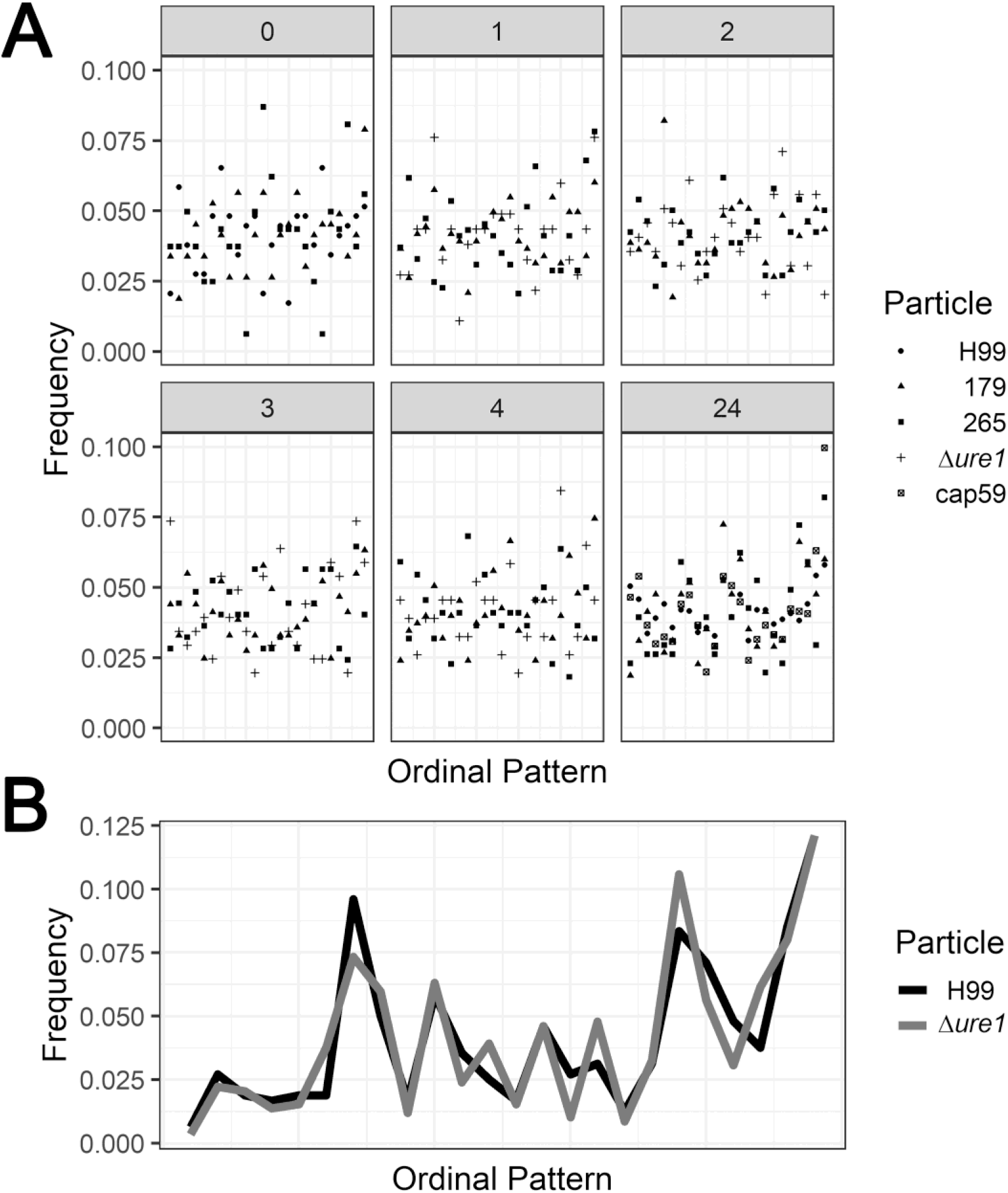
Ordinal pattern analysis for expanded samples. **A.** Ordinal pattern frequencies for phagolysosomal acidification intervals in BMDMs infected with various particles at various hours post infection (gray box values). **B.** Ordinal pattern frequencies for time elapsed before initiation of budding for yeast strains ingested by BMDMs.

**Supplemental Figure 3.**
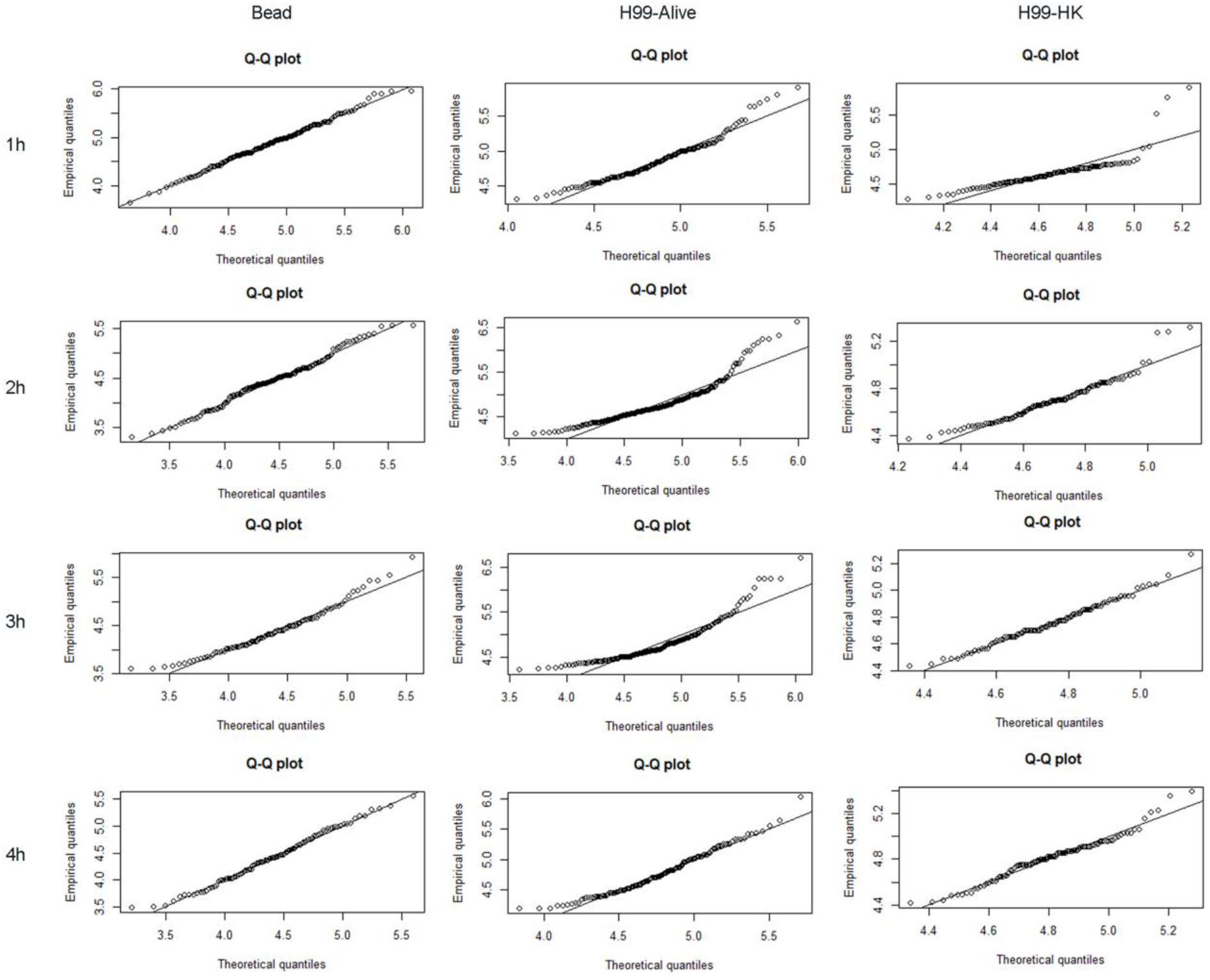
Q-Q plots of macrophages 1-4 HPI after ingesting various particles. Note that bead containing phagolysosomes closely approximate a normal distribution, while *C. neoformans* containing phagolysosomes have heavy tails, regardless if the particle is live or dead.

**Supplemental Figure 4.**
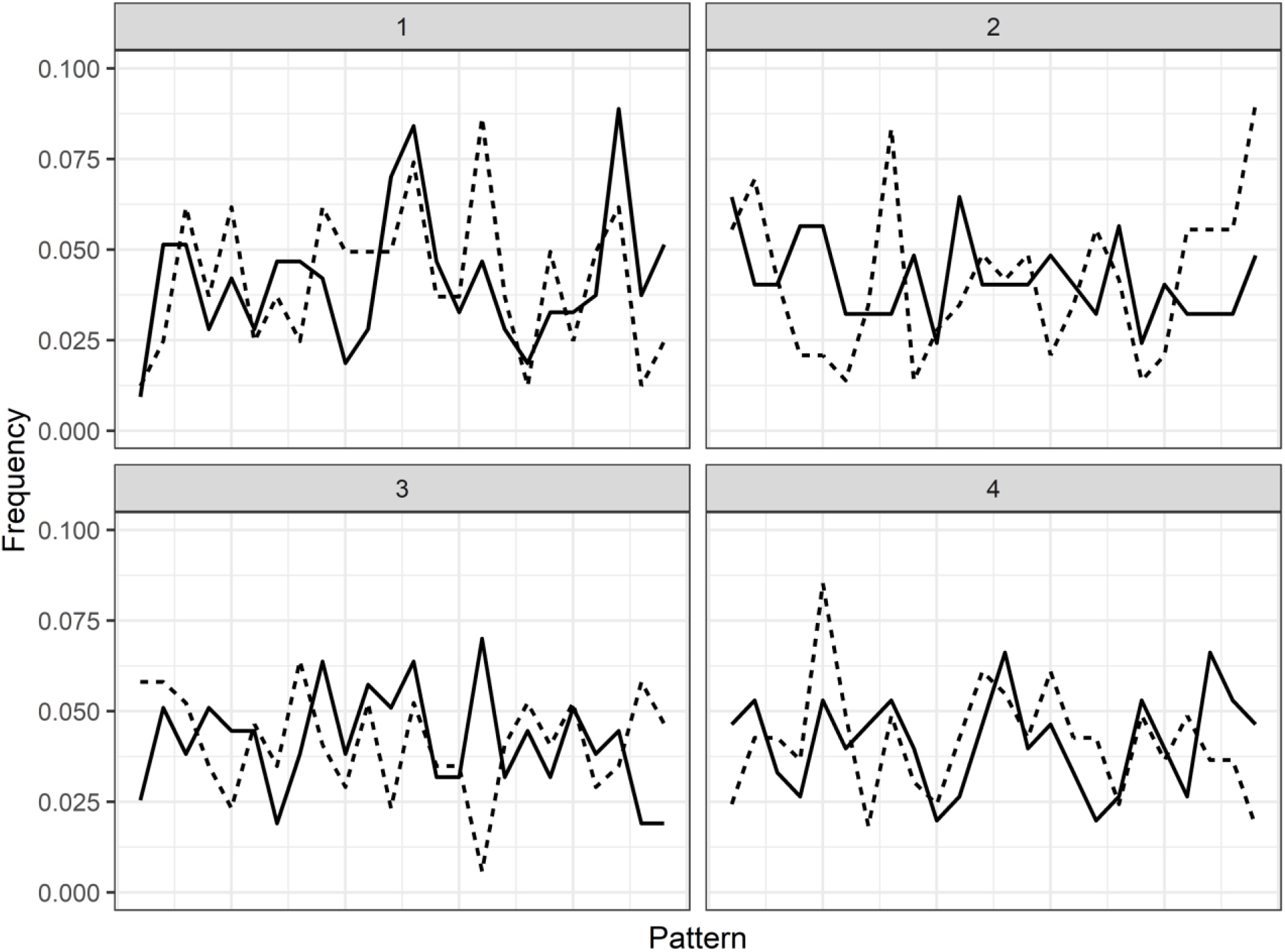
Ordinal pattern analysis for EEA1 (solid) and VATPase (dashed) immunofluorescent staining for bead containing macrophage phagolysosomes at various hours post ingestion (gray box values). No forbidden ordinal patterns were detected for any samples, suggesting phagolysosomal maturation marker acquisition is a stochastic process.

**Supplemental Figure 5.**
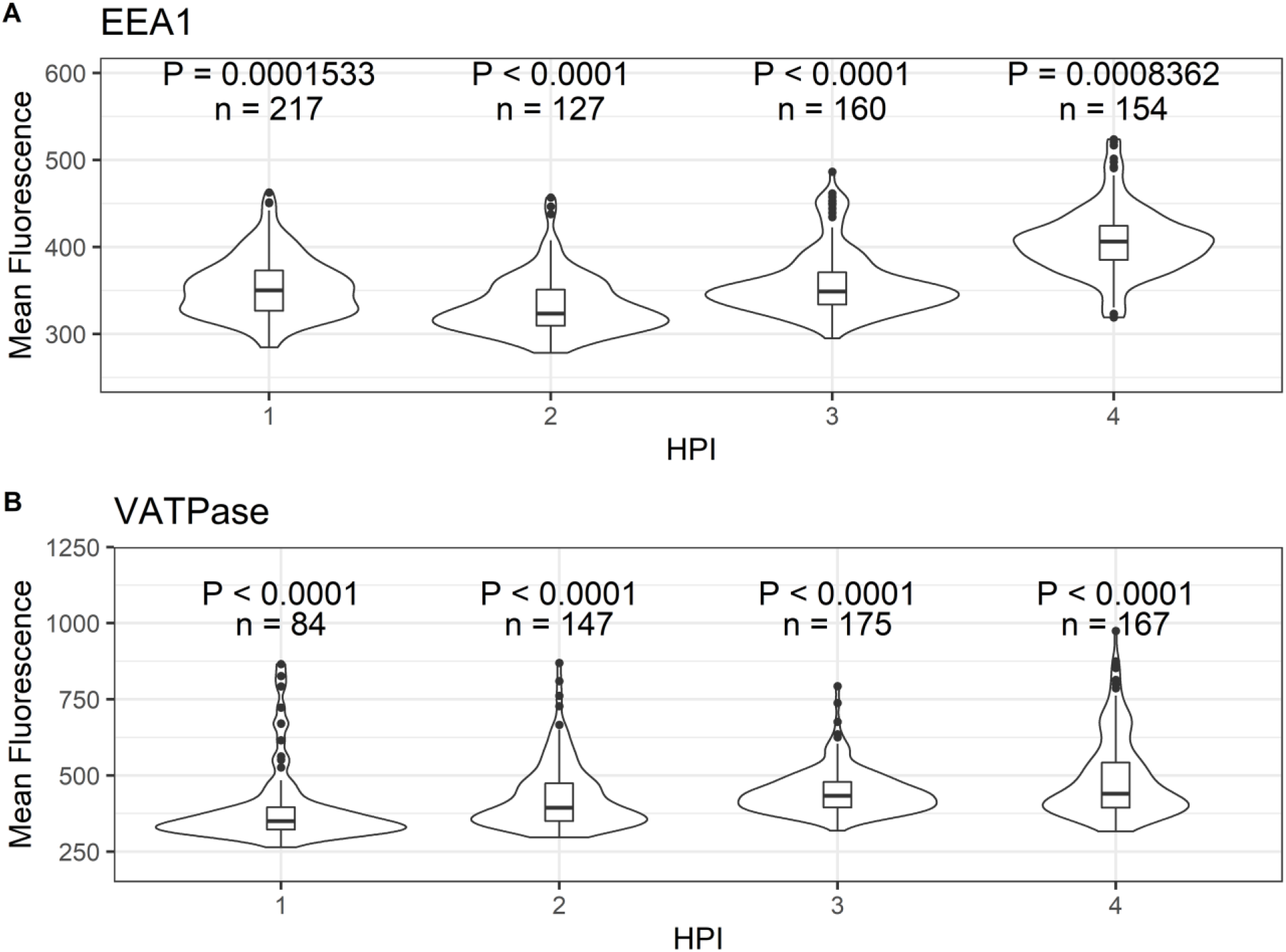
Mean fluorescence intensity values for EEA1 and VATPase immunofluorescent staining. The acquisition of these phagolysosomal maturation markers does not resemble that of bead containing phagolysosomal pH, as no samples here are normally distributed. P values were determined via Shapiro-Wilk normality test.

**Supplemental Figure 6.**
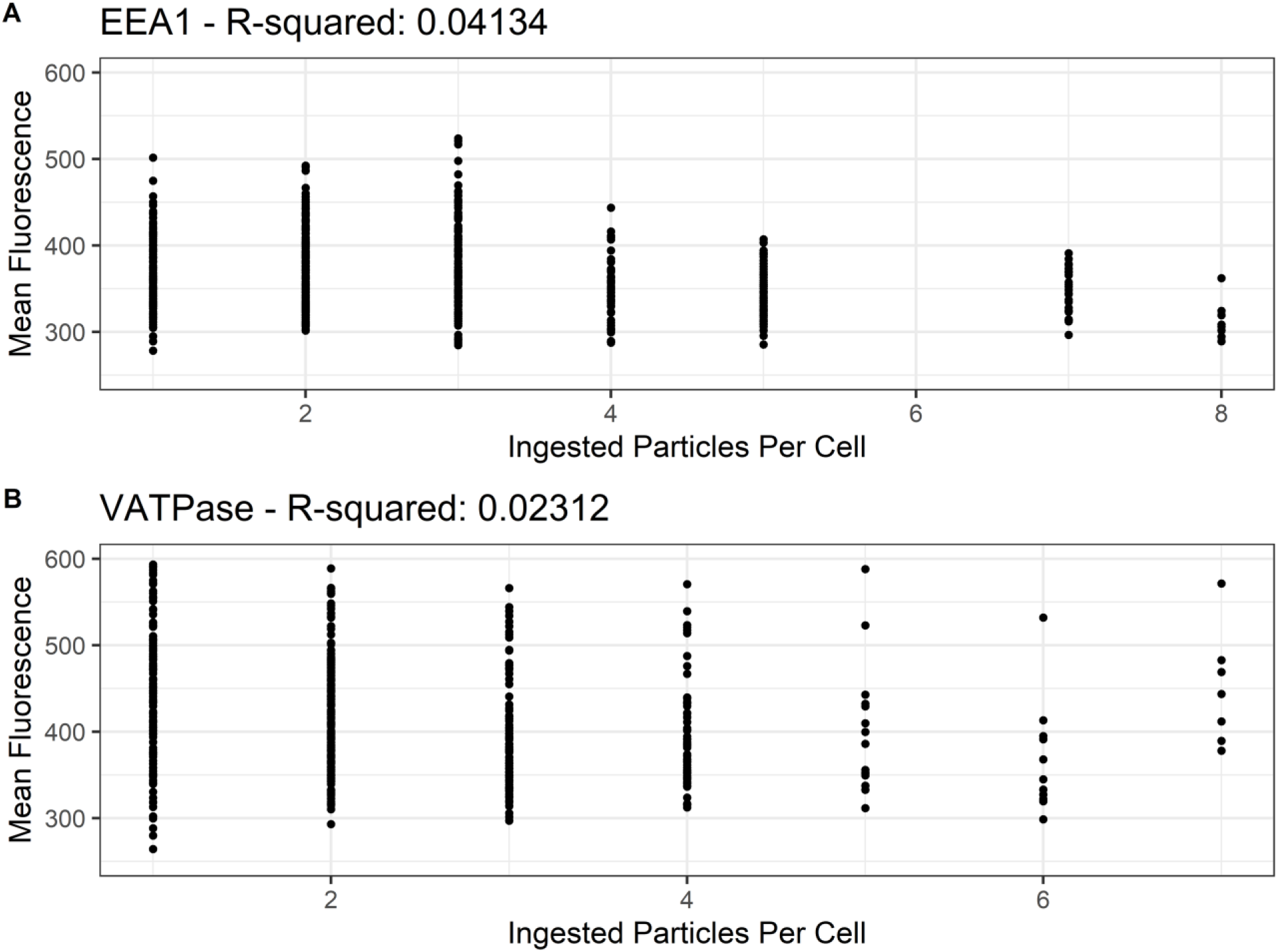
Mean fluorescent intensity of macrophage ingested beads stained for EEA1 and VATPase, sorted according to total ingested particles per cell. Linear regressions were calculated for each sample set but clearly there is no significant correlation between total ingested particles and number of EEA1 or VATPase molecules as measured by fluorescent intensity.

**Supplemental Figure 7.**
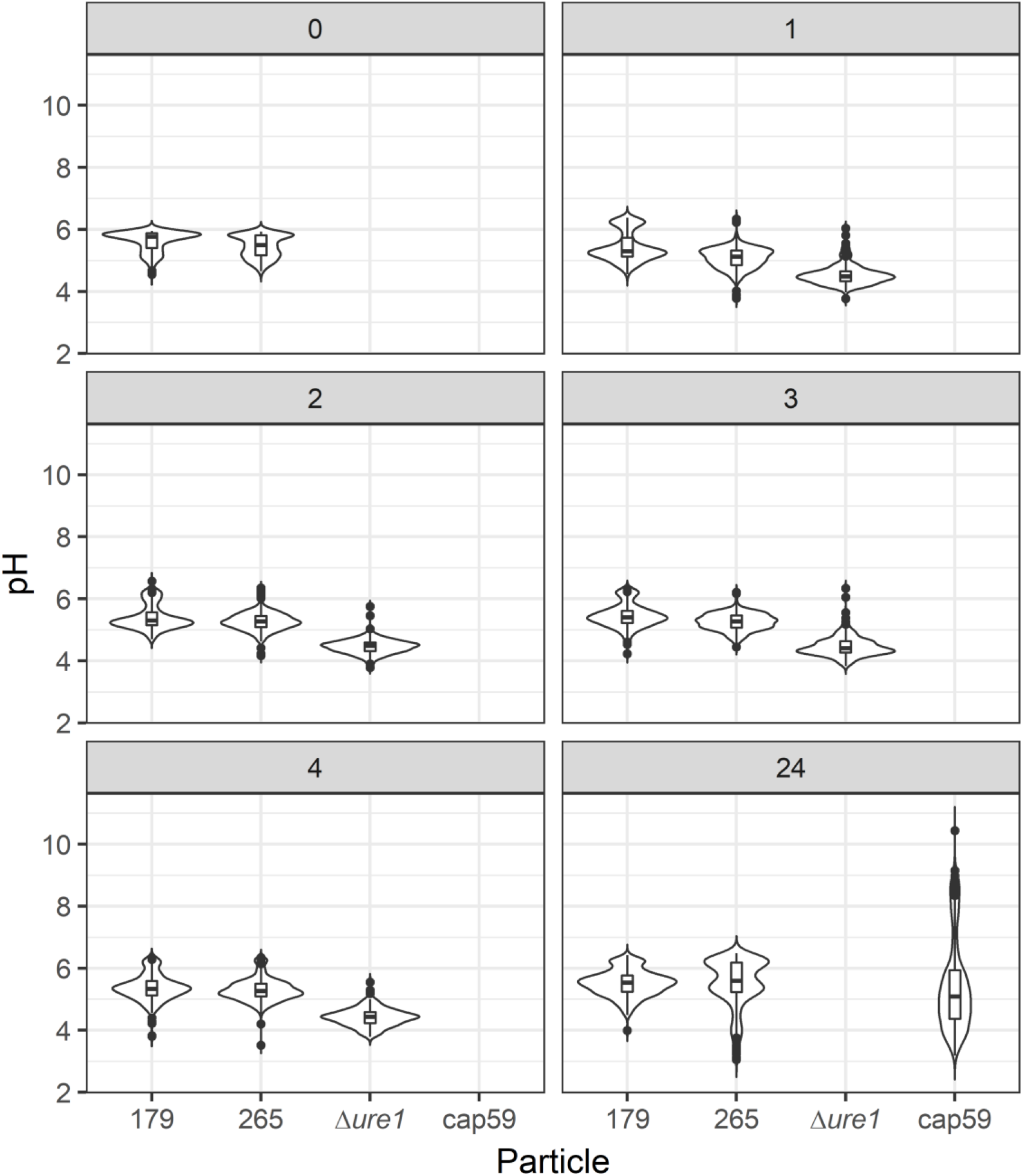
Phagolysosomal pH distributions for expanded samples. Two strains of *C. gattii* (179 and 265), a urease deficient H99 mutant (*Δure1),* and a capsule deficient H99 mutant (cap59) were analyzed at various hours post infection (gray box value).

**Supplemental Figure 8.**
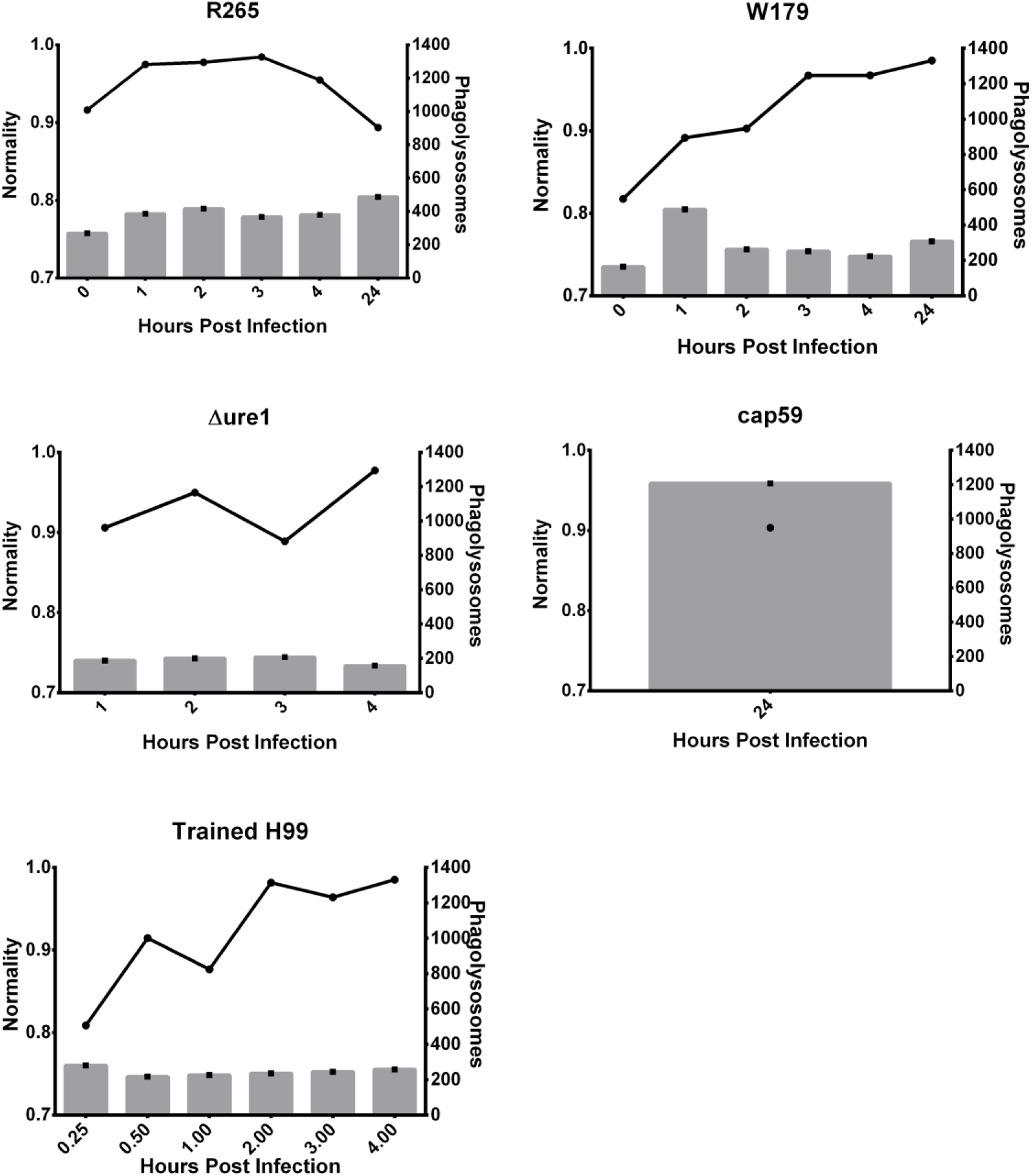
Normality analysis for expanded samples. Shapiro-Wilk normality values (lines) alongside total phagolysosome sample sizes (bars) for two strains of *C. gattii* (179 and 265), a urease deficient H99 mutant (*Δure1),* a capsule deficient H99 mutant (cap59), and trained BMDMs on reinfection.

**Supplemental Figure 9.**
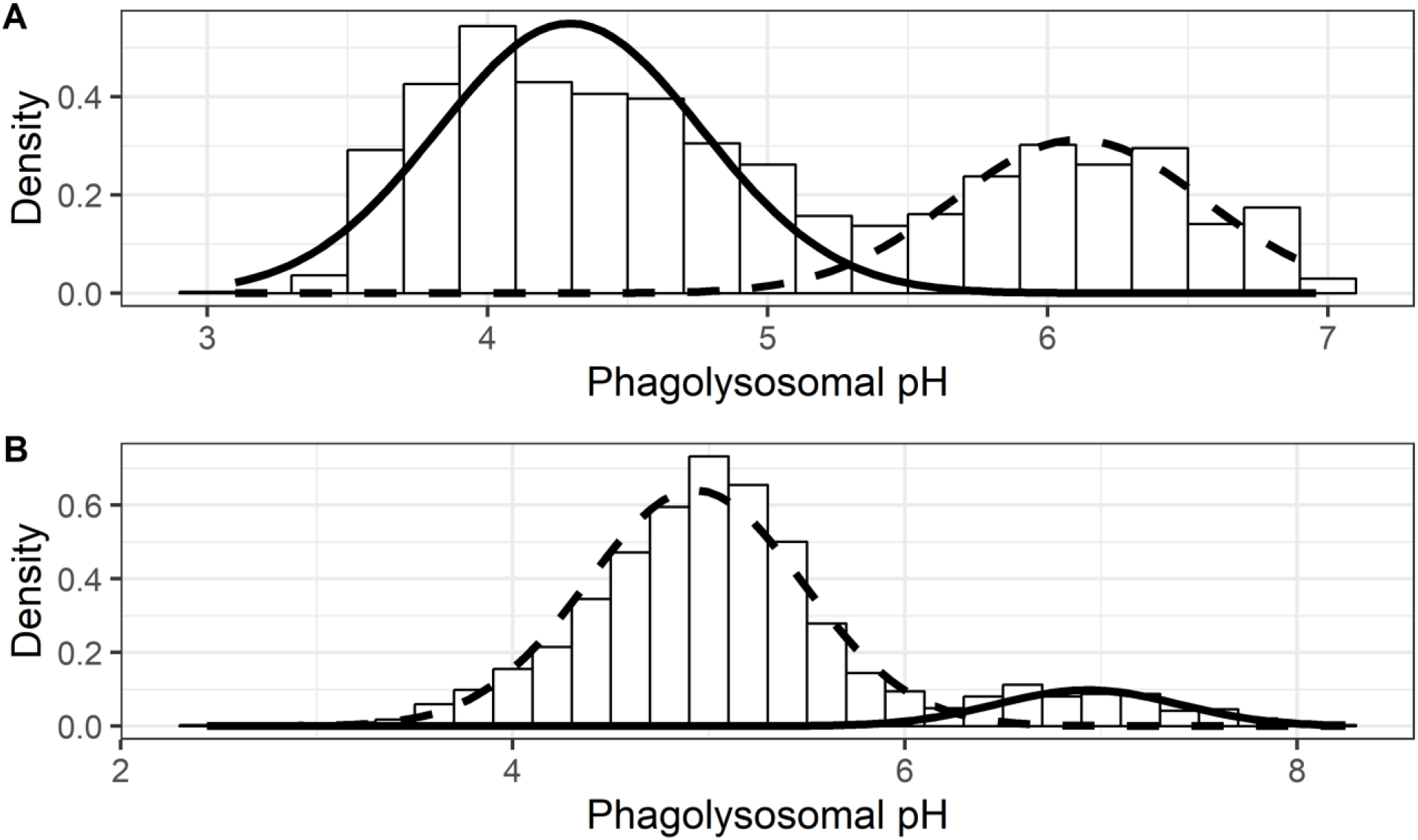
Bimodal models fitted to the observed data for **A.** M0 and **B.** M2 skewed macrophages. Models are attempted fits of two mixed Gaussian distributions. Histogram bars visualize the observed data while solid and dashed lines depict the relative contributions of the two hypothetical Gaussian distributions.

**Supplemental Figure 10.**
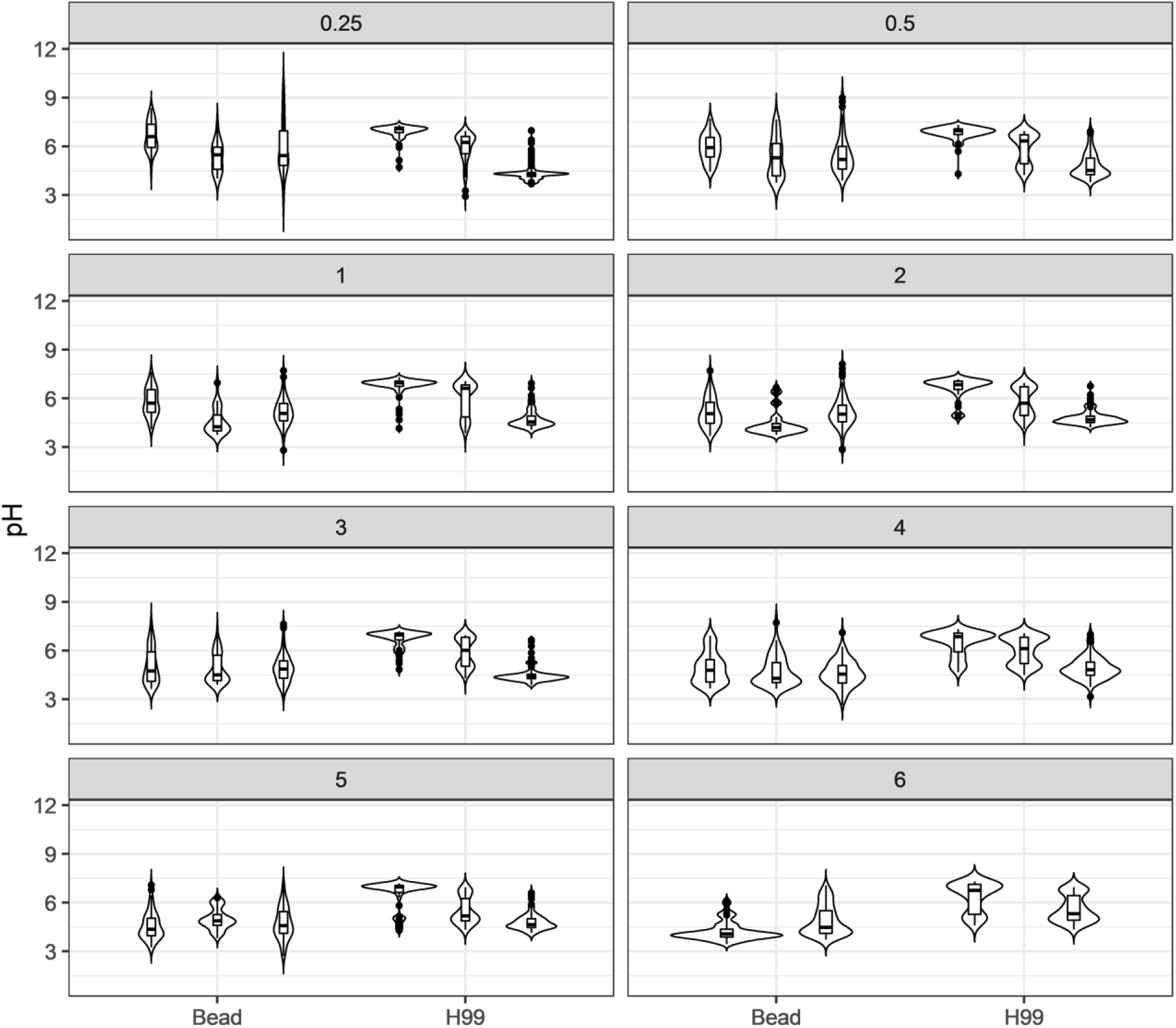
Phagolysosomal pH distributions for human macrophages infected with H99 separated by individual donor. Human macrophages were analyzed after ingesting either H99 or inert beads at various hours post infection (gray box value).

**Supplemental Figure 11.**
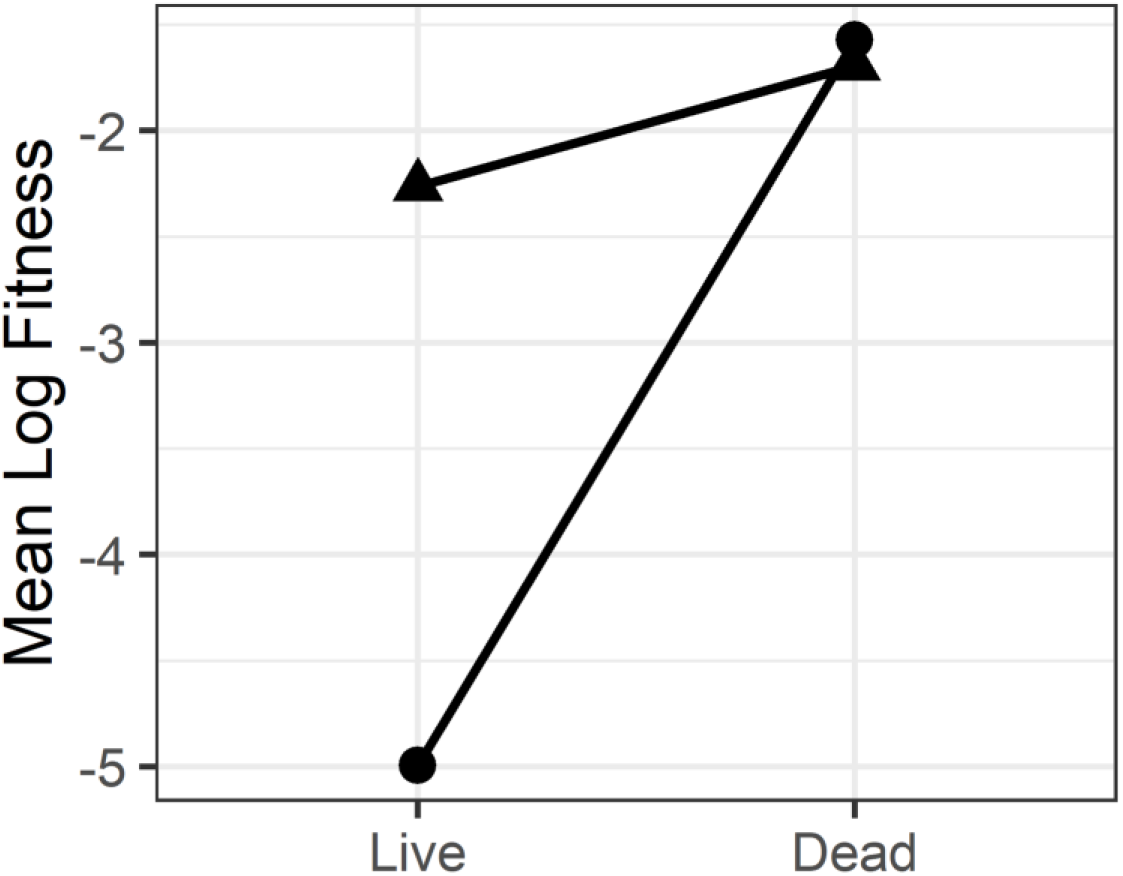
Estimated mean log fitness of macrophages containing live or killed *M. avium*. Phagolysosomal pH data was gathered from literature using either video microscopy (triangles) or confocal microscopy (circles).

## Notes

#### Summary of Updates

We have included many new experiments and insights as well as updated some analysis. The conclusions are unchanged but our methodology and writing have been improved.

